# Structures of AT8 and PHF1 Phospho-Mimetic Tau: Insights Into the Posttranslational Modification Code of Tau Amyloid Formation

**DOI:** 10.1101/2023.09.04.556256

**Authors:** Nadia El Mammeri, Aurelio J. Dregni, Pu Duan, Mei Hong

## Abstract

The microtubule-associated protein tau aggregates into amyloid fibrils in Alzheimer’s disease and other neurodegenerative diseases. In these tauopathies, tau is hyperphosphorylated, suggesting that this posttranslational modification may induce pathological tau aggregation. Tau is also phosphorylated in normal developing brains. To investigate how tau phosphorylation induces amyloid fibrils, here we report the atomic structures of two phospho-mimetic full-length tau fibrils assembled without anionic cofactors. One set of phospho-mimetic mutations is targeted by the antibody AT8, while the other set is targeted by the antibody PHF1. Solid-state NMR and cryo-electron microscopy data reveal that AT8 tau forms a unique triangular fibril core that encompasses the entire C-terminal third of the protein, whereas PHF1 tau forms a triple-stranded core. These results demonstrate that specific post-translational modifications induce structurally specific tau aggregates. We propose that these aggregates may evolve into pathological filaments under suitable cellular conditions or remain as transient species in normal brains.

## Introduction

The microtubule-associated protein tau aggregates into β-sheet rich amyloid fibrils in many neurodegenerative diseases ^1–3^. In Alzheimer’s disease (AD), the tau aggregates spread from the locus coeruleus to the entorhinal cortex, continue to the hippocampus, then the entire neocortex ^4–5^. This spatial spreading correlates with cognitive decline and is the basis for the neuropathological staging of AD. Cryo-electron microscopy (cryo-EM) structures of AD paired helical filament (PHF) tau ^6–9^ show the same structure between multiple individuals, both with and without bound small molecules that disaggregate or image the fibrils. Meanwhile, tau fibrils obtained from the brains of different tauopathies ^10–13^ show different structures. These results imply that specific tau conformations exist in the complex cellular milieu and are linked to specific pathology. Elucidating the chemical and environmental factors that drive the prion-like propagation and spreading of these tau conformations ^14–15^ is therefore essential for designing disease-specific diagnostic ligands and therapeutic compounds.

Tau aggregation is counter-intuitive, because native tau is highly charged, highly soluble, and resistant to aggregation ^16^. The electrostatic charges in native tau are segregated in its modular amino acid sequence: the long N-terminal domain (NT) and the short C-terminal domain (CT) are negatively charged at neutral pH, whereas the central portion of the protein, comprising a proline (Pro)-rich region (P1 and P2) and several microtubule-binding repeats (R), is positively charged. The microtubule-binding repeats are R1, R2, R3, R4 and R’ in 4R tau isoforms whereas 3R tau isoforms miss the R2 domain due to alternative splicing. In AD PHF tau, the negatively charged terminal domains form a disordered fuzzy coat around the positively charged rigid core ^17–18^. Full-length 0N4R tau, the most common isoform in adult human brains, has a net charge of +15 at neutral pH. Given this cationic nature, it is not surprising that tau interacts with many anionic cellular species, including microtubules ^19^, heparin ^20^, RNA ^21^, and lipid membrane ^22^. Similarly, charge-modifying post-translational modifications (PTMs) are common in tau. The most studied tau PTM is phosphorylation. The protein composition of AD PHF tau was identified based on its abnormal phosphorylation of S396 ^23^. About 45 phosphorylation sites have been found in insoluble AD PHF tau ^24^, most of which are Ser-Pro and Thr-Pro motifs in the regions flanking the repeats. Among these phosphorylation sites, two motifs are recognized by anti-phosphorylated tau antibodies: a motif involving S202, T205 and S208 in the P2 domain ^25–29^, recognized by the antibody AT8, and a motif involving S396, S400, T403 and S404 between the R’ domain and the CT, recognized by the antibody PHF1 ^30^. Indeed, AT8 is the most widely used antibody for detecting phosphorylated tau in AD and other tauopathies.

It is commonly hypothesized that abnormal phosphorylation, by reducing the positive charges, causes tau to detach from microtubules, thus inducing its self-assembly into pathological aggregates. Biochemical evidence that tau phosphorylation at certain sites weakens microtubule binding ^31^ and induces self-assembly ^29, 32^ have been reported. However, counter-evidence that phosphorylation at other sites protects against aggregation despite weaker microtubule binding has also been reported ^33^. Tau phosphorylation also occurs in human fetal brains and hibernating animals at some of the same sites as in tauopathy brains, but these phosphorylation events appear to be reversible ^34–36^. Given the overlapping patterns of tau phosphorylation in pathological and physiological conditions, structural information of phosphorylated tau assemblies can provide important clues to the question of how phosphorylation impacts tau aggregation.

To date, no high-resolution structures of full-length tau aggregates with phosphorylation or phospho-mimetic mutations have been reported. Solid-state NMR spectroscopy has been used to investigate the conformation and dynamics of unphosphorylated tau fibrils assembled using the polyanionic cofactor heparin ^37–43^. Fluorescence spectroscopy has been used to study the global fold of soluble tau that contains phospho-mimetic mutations at AT8 and PHF1 epitopes ^44^. Solution NMR has been used to study the local conformational dynamics of soluble tau with phospho-mimetic mutations ^29, 45^ or phosphorylation by kinases ^46–47^. Here we report the atomic structures of phospho-mimetic full-length 0N4R tau fibrils assembled without cofactors. We installed three Glu mutations at S202, T205 and S208 to create AT8-0N4R tau (abbreviated as AT8 tau) and four Glu mutations at S396, S400, T403, and S404 to create PHF1 tau. A third sample, AT8/PHF1 tau, contains all seven mutations. We show that the installation of only three and four negative charges are sufficient to induce homogeneous tau fibrils with a single predominant molecular structure. Combining solid-state NMR and cryo-EM, we determined the high-resolution structures of the rigid cores of these tau fibrils. By detecting the dynamic residues by NMR, we also obtained information about the dynamics and packing of the disordered flanking domains around the rigid core. These data indicate that the introduction of negative charges in the Pro-rich region allosterically exposes the most amyloidogenic repeat of the protein while sequestering the C-terminal domain. Introduction of negative charges at the R’-CT junction creates a three-stranded β-sheet core that is reminiscent of *ex vivo* fibrils of 4R tauopathies. These results provide a structurally based mechanistic framework for the development of hyperphosphorylated tau aggregates in human brains.

## Results

### AT8 and PHF1 tau form well-ordered aggregates in the absence of anionic cofactors

The amino acid sequence of 0N4R tau with the AT8 and PHF1 phospho-mimetic mutations are shown in **Fig. 1a**. We over-expressed the proteins in *E. coli* and purified them by heat denaturation, cation-exchange column chromatography, and HPLC. Fibrils were assembled from monomer solutions at 0.4 mg/mL for AT8 tau and AT8/PHF1 tau and at 1.6 mg/mL for PHF1 tau. The solutions were incubated at 37°C under shaking at 250 rpm for 14 days, during which DTT was added every 2 days to prevent cysteine oxidation. No cofactors were used. Fibrils began forming after one week, and the reactions were stopped after two weeks. Transmission electron micrographs revealed 200 nm long and 10-15 nm wide fibrils for the AT8 and PHF1 mutants (**Fig. 1b**). The cofactor-free fibrilization of these phospho-mimetic tau samples differs qualitatively from wild-type (WT) full-length tau, which does not form homogeneous fibrils without cofactors ^48^. Therefore, the introduction of only three or four negative charges is sufficient to induce self-assembly of full-length tau.

**Figure 1.**
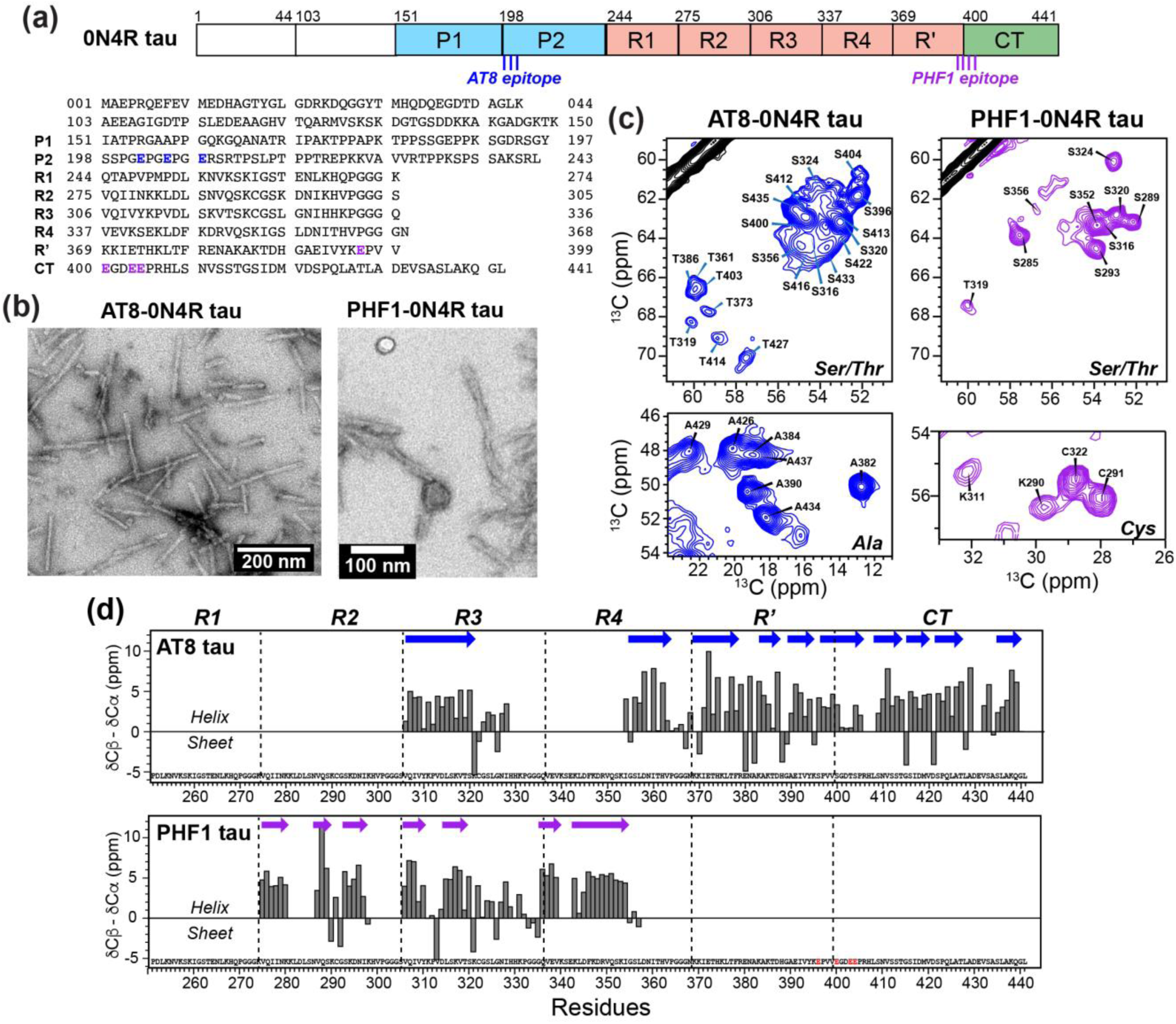
Amino acid sequences, fibril morphologies, and ssNMR spectra of full-length 0N4R tau fibrils containing phospho-mimetic mutations. (**a**) Domain diagram of 0N4R tau. The positions of the AT8-0N4R epitope and PHF1-0N4R epitope are indicated. The full amino acid sequence shows the AT8-0N4R mutations (S202E, T205E, S208E) and PHF1-0N4R mutations (S396E, S400E, T403E, and S404E). (**b**) Negative stain TEM images of full-length AT8-0N4R tau and PHF1-0N4R tau fibrils formed in the absence of any cofactors. These fibrils were used for ssNMR and cryo-EM characterization. (**c**) Representative regions of the 2D dipolar ^13^C-^13^C correlation spectra of the two phospho-mimetic tau fibrils. The Ser/Thr region and the Ala and Cys regions are shown. A single set of ^13^C chemical shifts is observed in each fibril, indicating that both AT8-0N4R and PHF1-0N4R 0N4R tau fibrils adopt single molecular conformations. (**d**) Secondary-structure dependent chemical shifts of the rigid cores of AT8-0N4R– and PHF1-0N4R tau fibrils. The difference between the secondary chemical shifts of Cβ and Cα (δCβ – δCα) are plotted. Positive differences indicate β-strands while negative differences indicate coil or helical conformations.

### AT8 and PHF1 tau fibrils have different β-sheet conformations

To determine the position of the rigid core in the amino acid sequence of AT8 and PHF1 tau fibrils and characterize their secondary structures, we measured 2D and 3D correlation magic-angle-spinning (MAS) NMR spectra. 2D ^13^C-^13^C (CC) and NCACX correlation spectra provide the fingerprints of the rigid cores. Both AT8 and PHF1 fibrils show well-resolved spectra (**Fig. S1, S2**), indicating that the rigid cores are well ordered. The 2D CC spectra exhibit fewer than 20 Ser and Thr peaks for AT8 tau and fewer than 10 peaks for PHF1 tau (**Fig. 1c**), suggesting that AT8 tau has a larger fibril core. To assign the ^13^C and ^15^N chemical shifts, we measured 3D NCACX, NCOCX, and CONCA correlation spectra (**Fig. S3**). For AT8 tau, we assigned 106 residues from the beginning of R3 (V306) to the C-terminus (G440), except for residues H330 to I354, which do not show signals in these dipolar NMR spectra, indicating they are dynamically disordered (**Fig. 1d, Table S1**). The assigned chemical shifts indicate the existence of ten β-strands, separated by Pro and Gly-rich segments such as ^322^CGS^324, 364^PGGG^366^, and ^389^GAE^391^. The caspase cleavage site D421 shows non-β-strand chemical shifts and acts as the break between the eighth and ninth β-strands. For PHF1 tau, we assigned 68 residues from the beginning of R2 (V275) to the middle of R4 (L357) (**Table S2**). This rigid core contains seven identifiable β-strands. For both fibrils, most strong peaks are assigned, and a single set of chemical shifts are observed, indicating that both fibrils adopt a single predominant molecular conformation. Strikingly, when Glu mutations are introduced to both AT8 and PHF1 sites, the protein shows an almost identical 2D NCA spectrum as PHF1 tau (**Fig. S4**), indicating that the seven-Glu tau has a very similar amyloid structure as the four-Glu PHF1 tau.

### Fibril core structures of AT8 tau and PHF1 tau

With the rigid cores identified by NMR chemical shifts, we next turned to cryo-EM to obtain the three-dimensional folds of these phospho-mimetic tau fibrils. Because the AT8/PHF1 tau has the same NMR spectra as PHF1 tau, we focused on the two primary mutants for structure determination. Fibrils were manually picked from about 11,000 micrographs for each fibril sample (**Fig. S5, Fig. 2**). Manual alignment of the resulting 2D classes led to an initial estimate of the crossover length to be ∼80 nm for PHF1 tau and ∼150 nm for AT8 tau. This crossover length estimate was crucial for successful 3D classification and refinement, which resulted in a 2.6 Å resolution cryo-EM map for AT8 tau and a 2.4 Å cryo-EM map for PHF1 tau (**Table 1**). The AT8 tau fibril core has a complex multilayered triangular shape whereas the PHF1 tau fibril core exhibits a simple three-layered β-strand fold (**Fig. 3**). A single structural model was found for each fibril, consistent with the observation of a single set of NMR chemical shifts. For both samples, the structures comprised one protofilament, with a refined helical crossover distance of 123 nm for AT8 tau and 79 nm for PHF1 tau (**Fig. 2**).

**Figure 2.**
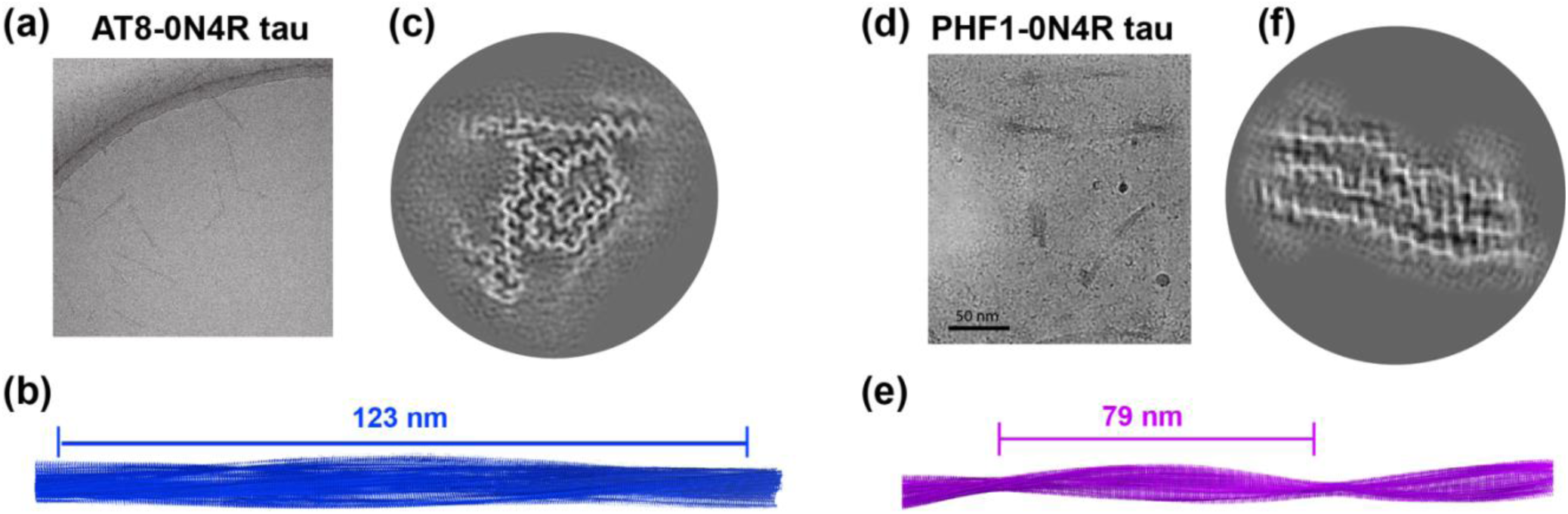
Cryo-EM data of two phospho-mimetic full-length 0N4R tau. (**a-c**) AT8 tau fibril data. (**a**) Representative micrograph. (**b**) Helical reconstruction of AT8 tau fibrils with a crossover length of 123 nm. (**c**) Projected slice of the central 4.8 Å of the final 3D reconstructed map of AT8 tau. (**d-f**) PHF1 tau fibril data. (**d**) Representative micrograph. (**e**) Helical reconstruction of PHF1 tau fibrils with a crossover length of 79 nm. (**f**) Projected slice of the central 4.8 Å of the final 3D reconstructed map.

**Figure 3.**
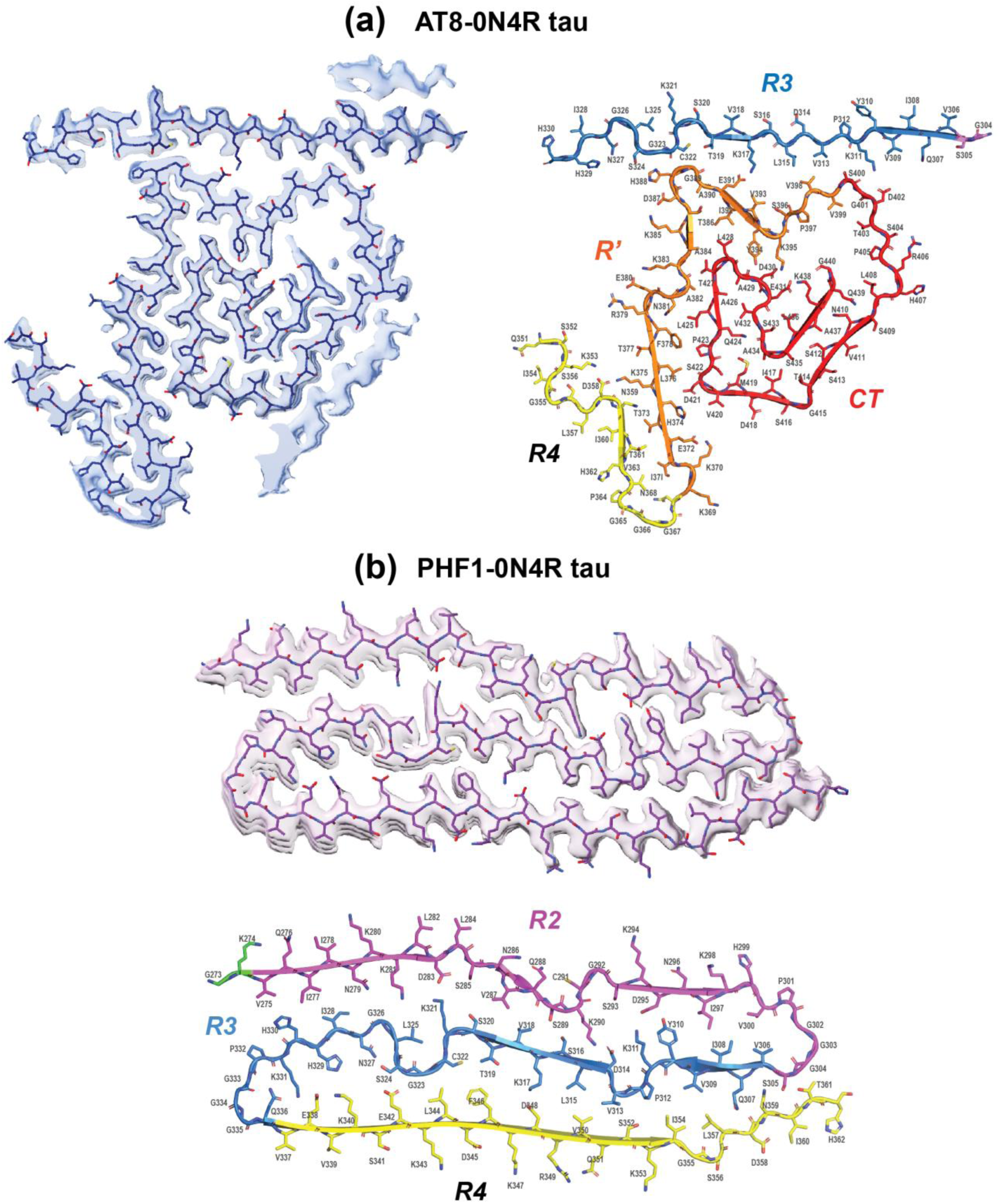
High-resolution structures of the rigid cores of phospho-mimetic 0N4R tau fibrils. (**a**) The AT8 tau rigid core adopts a complex triangular structure that spans the R3 to the CT, except for the last 7 residues of R3 and the first half of the R4, which are disordered. (**b**) The PHF1 tau rigid core spans R2, R3 and R4, which form antiparallel stacked β-strands that are separated by PGGG motifs.

**Table 1.** Refinement statistics for AT8-0N4R and PHF1-0N4R 0N4R tau fibrils.

|  | AT8-0N4R tau<br>EMD 41610,<br>PDB 8TTL | PHF1-0N4R tau<br>EMD 41611<br>PDB 8TTN |
| --- | --- | --- |
| **Data collection and processing** |  |  |
| Microscope | Krios G3i | Krios G3i |
| Camera | Bioquantum K3 | Bioquantum K3 |
| Energy Filter (eV) | 20 | 20 |
| Magnification | 130,000 | 130,000 |
| Voltage (kV) | 300 | 300 |
| Electron exposure (e<sup>-</sup>/ Å<sup>2</sup>) | 48.7 | 50.7 |
| Defocus Range (µm) | -1.75 to 0 | -2 to -0.4 |
| Pixel Size (Å) | 0.654 | 0.654 |
| Initial no. particle images | 675,445 | 3,830,683 |
| Final no. particle images | 21,767 | 132,972 |
| Helical twist (°) | -0.7 | -1.07 |
| Helical rise (Å) | 4.80 | 4.79 |
| Symmetry imposed | C1 | C1 |
| Map resolution FSC 0.143 (Å) | 2.6 | 2.4 |
| FSC threshold | 0.143 | 0.143 |
| **Refinement** |  |  |
| Initial model used (PDB code) | - | - |
| Model resolution (Å) | 3.1 | 3.14 |
| FSC threshold | 0.5 | 0.5 |
| Map sharpening *B* factor (Å<sup>2</sup>) | -28.18 | -23.67 |
| Model composition |  |  |
| Non-hydrogen atoms | 2583 | 3380 |
| Protein residues | 351 | 450 |
| Ligands | - | - |
| *B* factors (Å<sup>2</sup>) |  |  |
| Protein | 48.8 | 68.2 |
| Ligand | - | - |
| R.m.s. deviations |  |  |
| Bond lengths (Å) | 0.011 | 0.012 |
| Bond angles (°) | 2.141 | 2.556 |
| Validation |  |  |
| MolProbity score | 1.31 | 1.24 |
| Clashscore | 0.00 | 1.01 |
| Poor rotamers (%) | 2.36 | 0.51 |
| Ramachandran plot |  |  |
| Favored (%) | 90.86 | 93.18 |
| Allowed (%) | 8.26 | 4.32 |
| Disallowed (%) | 0.88 | 2.50 |

Based on the chemical-shift derived β-strand positions and the cryo-EM densities, we generated atomic models of the two phospho-mimetic tau fibrils. Out of 383 residues of 0N4R tau, AT8 tau comprises 115 residues in its rigid core, spanning residues K305-H330 and Q351-G440 (**Fig. 3a**). The discontinuity between H330 and Q351 is in excellent agreement with the lack of signals for these residues in the dipolar NMR spectra (**Fig. 1d**). Thus, the 3% of the particles used for cryo-EM helical reconstruction are consistent with dipolar NMR spectra that detect all rigid proteins. The triangular shape of AT8 tau’s rigid core is bounded by the first 25 residues of R3 (V306-H330) at the top, the second half of R4 (Q351-G367) on the left, and the first half of CT (R406-S416) on the right. This triangle is filled by four layers of β-strands in the interior. After the R4 ^364^PGGG^367^ segment on the left side of the triangle, the protein makes a sharp U-turn, packing the first half of R’ against the second half of R4. This R’ β-strand continues until it reaches the ^322^CGS^324^ motif in the R3 strand, then makes a sharp right turn at H388 to stack the second half of R’ (H388-S400) against the R3 strand. This antiparallel R’-R3 stacking is stabilized by hydrophobic interactions between L315 and V398, salt bridge between K317 and E391, and a second salt bridge between K311 and D402 (**Fig. S6b**). Notably, K311 is part of the amyloidogenic R3 hexapeptide whereas D402 is part of the PHF1 epitope. C-terminal to D402, the protein spirals inward, turning sharply at H407, G415, D421, L428, E431 and S435. The chemical shifts of these residues are consistent with the non-β-strand conformations seen in the cryo-EM densities.

PHF1 tau adopts a simpler β-sheet structure than AT8 tau: the triple-stranded core is bookended by the PGGG motifs at the end of the R2 and R3 repeats (**Fig. 3b**). The outer layers are composed of the R2 residues K274-P301 and R4 residues V337-H362. R3 forms the central strand and establishes numerous sidechain contacts with R2 and R4 strands. The antiparallel R3-R2 interface is stabilized by hydrophobic interactions such as I308-I297, polar interactions such as S316-S289, and salt-bridge interactions at K311-D295, D314-K290, and K321-D283. The antiparallel R3-R4 interface is more polar, stabilized by sidechain pairs such as K317-D348, Q307-N359, and S305-T361. The three β-strands are nearly perfectly matched with the three repeats, with the terminal PGGG in each repeat forming U-turns between the layers.

Between these two entirely different three-dimensional structures of AT8 and PHF1 tau, the R3 repeats adopts an uncannily similar conformation (**Fig. S6a**). In both fibrils, the ^320^SKCGSLG^3^^26^ segment forms an inverted-Ω fold, with the L325 sidechain filling the space within the loop. One notable difference in the R3 structures of the two fibrils is K311. In PHF1 tau, the K311 sidechain points to the same side of the β-strand as Y310, leading to a bent backbone that is consistent with the non-β-strand chemical shifts of ^311^KPV^313^. This bent backbone allows K311 to form a salt bridge with D295 in R2. In AT8 tau, the K311 sidechain lies on the opposite side of Y310 and P312, consistent with a canonical β-strand and the NMR chemical shifts of these residues. With this orientation, K311 forms a putative salt bridge with D402.

### Water pockets exist inside the fibril cores

To investigate whether water pockets exist inside the rigid cores, we measured water-edited 2D CC and NCACB correlation spectra (**Fig. 4, Fig. S7**). By transferring water ^1^H magnetization to the protein and detecting it through protein ^13^C signals, we probe the water accessibilities of individual residues. For AT8 tau, G440, V399, P397, and T403 are better hydrated than their neighboring residues, indicating the presence of a small water pocket that hydrates the C-terminus (**Fig. 4c**). L441 is not observed in either dipolar NMR spectra or cryo-EM densities, indicating that this C-terminal residue is disordered. On the left side of the triangular core, R’ residues K369-H388 are well hydrated, suggesting the presence of a water channel bounded by the disordered R4 repeat. At the top of the triangle, residues I308 and Y310 are water-inaccessible, suggesting that the R3 hexapeptide is shielded by the unidentified densities (**Fig. 3a**). In comparison, the H407-S416 stretch on the right side of the triangle is relatively water-accessible. This suggests that the parallel ridge of unidentified densities outside this segment is only present in some of the fibrils. Model building using Model Angelo ^49^ suggests that these unidentified densities may correspond to R2 residues K281-Q288, with a correlation coefficient of 0.3 between the map and the model (**Fig. S8**). Since solid-state NMR spectra report the water accessibility of all proteins whereas the cryo-EM structural model reflects a small percentage of all fibrils, these extra densities may reflect a subset of samples in which R2 is ordered and packed against the H407-S416 segment. Consistent with this interpretation, the N410-S416 segment has an average water-edited intensity ratio of ∼0.25 whereas the R3 hexapeptide has an average water-edited intensity ratio of less than 0.2. These indicate that the R3 hexapeptide is more shielded from water by the extra densities compared to the CT segment H407-S416.

**Figure 4.**
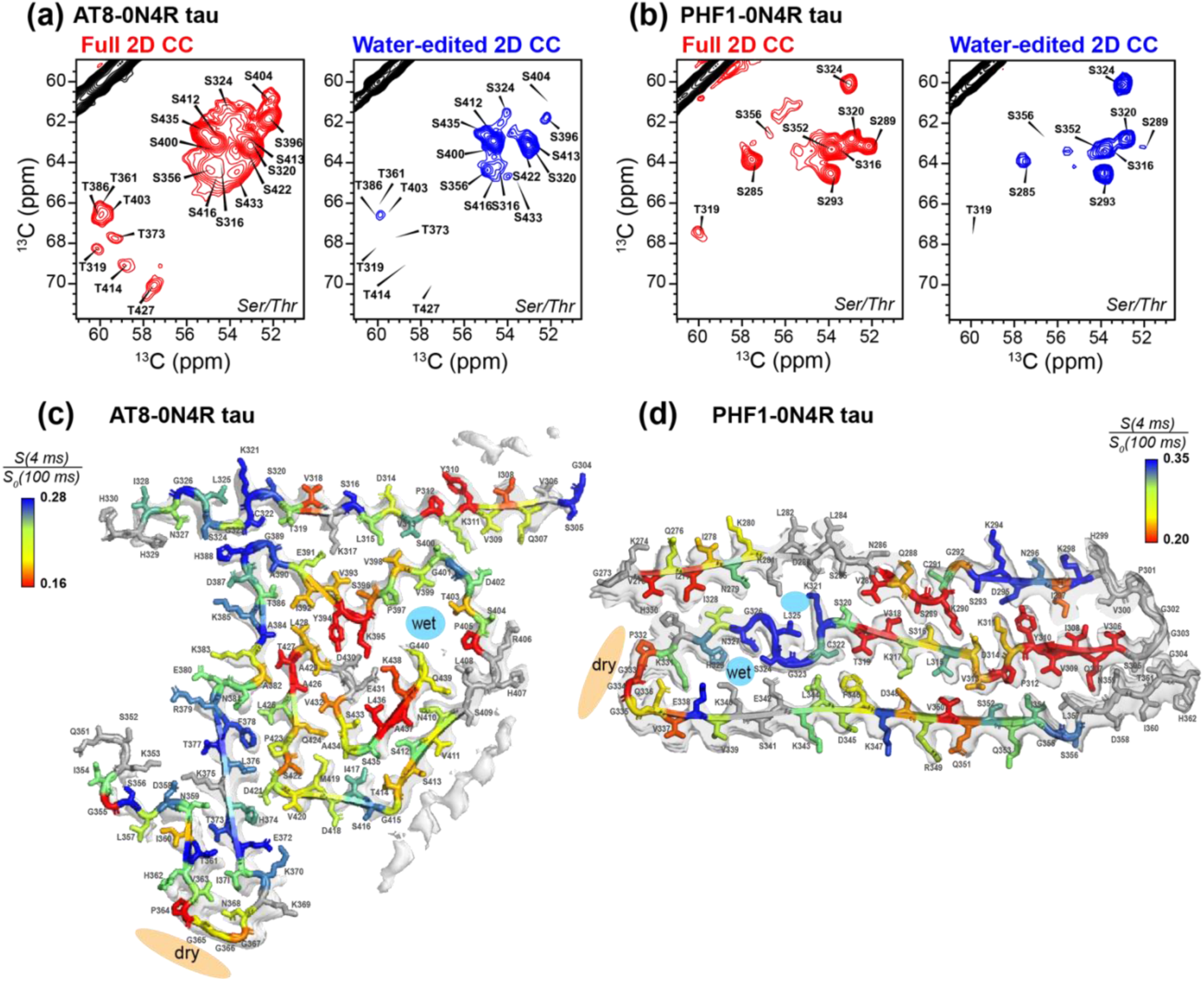
Dynamics and water accessibility of the phospho-mimetic tau fibrils. (**a**) Regions of full (red) and water-edited (blue) 2D CC spectra of AT8 tau fibrils. (**b**) Regions of full (red) and water-edited (blue) 2D CC spectra of PHF1 tau fibrils. Substantial intensity variations in the water-edited spectra are observed, indicating that the rigid core has heterogeneous water accessibilities. Low-intensity residues are less water accessible. (**c**) Residue-specific water accessibilities of AT8 tau fibrils. (**d**) Residue-specific water accessibilities of PHF1 tau fibrils. Hydrophobic sidechains in steric zippers are generally poorly accessible to water whereas surface-exposed residues are well hydrated. However, some internal hydrophobic residues are water accessible, suggesting the presence of internal water pockets. Some surface-exposed residues are dehydrated, suggesting that they are shielded from solvent by disordered regions of the protein.

The PHF1 tau fibril exhibits low water accessibility for internal hydrophobic steric zippers such as ^306^VQIVY^310^ but high water accessibility for surface residues such as ^293^SKDN^296^ (**Fig. 4d**). Interestingly, the inverted Ω motif ^320^SKCGSLG^326^ is well hydrated even though they are packed on both sides by R2 and R4 residues. This suggests the presence of two water pockets, one between R2 and R3 and bounded by K281, D283, K321 and L325, and the other between R3 and R4 and bounded by S324, H329, E338, K340, and E342. These water pockets solvate the hydrophilic sidechains in the interior of this fibril. In contrast, many surface-facing hydrophobic residues such as ^332^PGGGQV^337^ and the R2 hexapeptide ^275^VQIINK^280^ are water-inaccessible, suggesting that disordered segments are present to shield these hydrophobic residues from water exposure (*vide infra*).

### The three phospho-mimetic tau fibrils have different fuzzy coat dynamics

The dynamically disordered fuzzy coat of tau fibrils is invisible in dipolar NMR spectra and cryo-EM densities, but is important for mediating the interactions of the rigid β-sheet core with small molecules and other proteins ^50^. The fuzzy coat of tau also contains the majority of known phosphorylation sites, including AT8 and PHF1. Thus it is important to investigate the dynamics and shape of the fuzzy coat and their perturbation by phosphorylation. We measured scalar-coupling based 2D ^13^C-^13^C TOCSY and ^1^H-^15^N INEPT spectra under MAS to selectively detect highly mobile residues in the three tau fibrils (**Fig. 5, Fig. S9**). All three fibrils exhibit many sharp signals at approximately random coil chemical shifts, indicating the presence of highly dynamic residues. The ^13^C-^13^C TOCSY spectra resolved the signals of glutamate residues that precede proline (Glu-Pro) from those that do not (Glu). Interestingly, the relative intensities of these Glu-Pro and Glu peaks differ among the three samples: AT8 tau has a higher Glu-Pro intensity than Glu, whereas the other two fibrils have weaker Glu-Pro intensities than Glu. Other pairs of residues, including Ser-Pro versus Ser, Thr-Pro versus Thr, and Ala-Pro versus Ala, also show different intensity ratios among the three fibrils. Similarly, 2D ^15^N-^1^H INEPT spectra resolved many N-terminal residues, whose intensities show structurally informative variations among the three fibrils. AT8/PHF1 tau has the highest intensities for the N-terminal residues, whereas AT8 tau has the weakest intensities, indicating the N-terminal domain is partly immobilized in AT8 tau. Combining these TOCSY and INEPT spectral intensity patterns (**Fig. S9, Table S3**), we conclude that the NT and R1 are partially immobilized in AT8 tau, whereas the NT, R’ and CT are all partially immobilized in PHF1 tau (**Fig. 5c**). The AT8/PHF1 tau contains the most dynamic NT among the three fibrils while its R’ and CT are partially immobilized. These results are schematically depicted in **Fig. 5d**. We hypothesize that the partly immobilized NT in AT8 tau lies close to the R3 domain, reminiscent of the MC-1 antibody epitope ^51^. In comparison, the PHF1 and AT8/PHF1 fibrils contain a dynamic NT that resides further away from the rigid core in which the R3 is sequestered in the interior.

**Figure 5.**
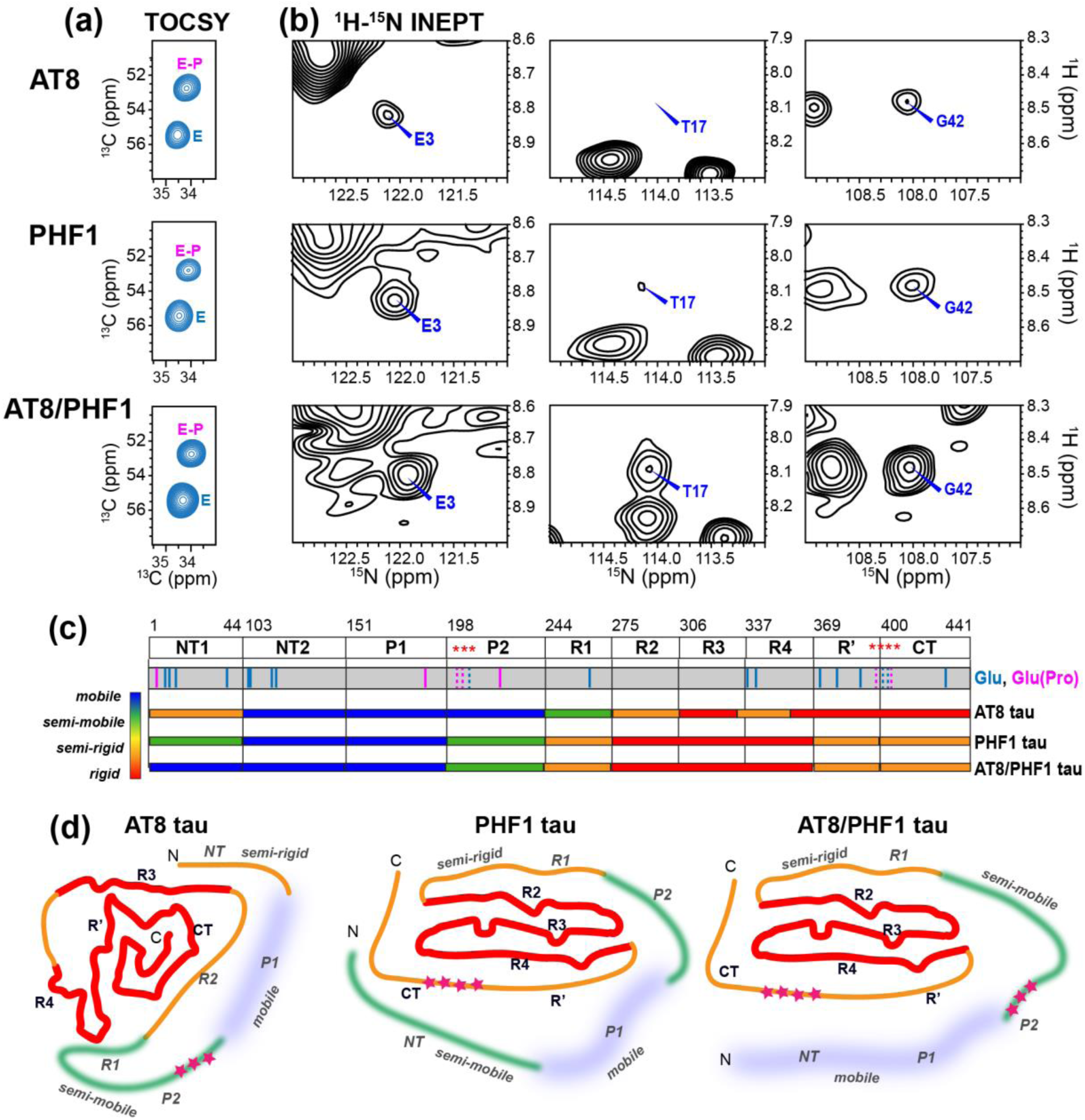
The fuzzy coat of the three phospho-mimetic tau fibrils has different dynamics. (**a**) Glu region of the 2D ^13^C-^13^C TOCSY spectra of AT8, PHF1, and AT8/PHF1 tau fibrils. For AT8 tau, the Glu-Pro (E-P) peak is more intense than the Glu (E) peak in the TOCSY spectrum. But for the other two fibrils, the E-P peak is weaker than the E peak. (**b**) Diagnostic regions of the 2D ^15^N-^1^H INEPT spectra of the three fibrils, showing distinct intensity distributions of N-terminal E3, T17 and G43 among the three samples. These intensity variations suggest that AT8/PHF1 tau has a highly dynamic NT, PHF1 tau is partially immobilized at T17 but retains a highly dynamic E3 and G42, whereas AT8 tau is less mobile at all three residues. (**c**) Distribution of Glu (blue lines) and Glu-Pro (pink lines) residues in the amino acid sequence of 0N4R tau, and summary of the mobilities of the three phospho-mimetic tau fibrils based on the NMR spectra. (**d**) Proposed arrangement of the fuzzy coat around the rigid core based on the NMR spectral intensities. These fuzzy coat models take into account the NT/CT interaction observed from FRET and solution NMR ^44–45^.

## Discussion

### Distinct tau phosphorylation patterns have distinct amyloid structures and the phosphorylation code contains redundancy and dominance

The high-resolution structures obtained here show that the addition of a small number of negative charges are sufficient to cause the highly soluble tau to aggregate into specific and distinct structures without cofactors. The introduction of three negative charges in the P1-P2 junction generated a large triangular β-sheet core that spans the segment from R3 to CT, except for ∼20 residues between R3 and R4. Thus, reduction of positive charges in the Pro-rich region allosterically sequestered R’, the highest-affinity MT-binding anchor of tau ^52^, while simultaneously exposing the most amyloidogenic segment of the protein, R3. The introduction of four negative charges at the R’-CT junction caused a three-layered β-sheet core that is evenly divided among R2, R3 and R4 repeats and that shows substantial similarities with a number of brain tau structures (*vide infra*). Thus, the native tau sequence is finely balanced between intrinsic disorder and order, and even small electrostatic changes can tip the balance towards aggregation.

The qualitatively different structures of AT8 tau and PHF1 tau indicate that distinct phosphorylation patterns lead to distinct amyloid structures. But the near identity of the NMR spectra of PHF1 tau and AT8/PHF1 tau (**Fig. S4**) implies that the phosphorylation code of tau has redundancy. When both AT8 and PHF1 sites are pseudo-phosphorylated, the resulting fibril adopts the same conformation as when PHF1 sites alone are pseudo-phosphorylated. Thus, the unique structural effects of AT8 phosphorylation are abolished by PHF1 phosphorylation, and the three-layered structure of PHF1 tau dominates the triangular structure of AT8 tau. This dominance may occur because specific interactions involving the AT8 sites are overridden by those of PHF1, or because PHF1-driven nucleation and elongation are faster than those of AT8. We can rationalize the first scenario by the similar R3 conformation in the two structures (**Fig. S6**). This R3 conformation differs mainly at K311, which forms a putative salt bridge with D402 in AT8 tau, which is absent in PHF1 tau. Thus, if monomers containing PHF1 phosphorylation near D402 are added onto a seed of AT8-like structure, the K311-D402 salt bridge in the seed structure would be disrupted, preventing the CT from interacting with the R3. This may then allow the R3 segment to scaffold against R2 and R4, potentially leading to the three-layered PHF1 fold (**Fig. 6b**). The identity of the PHF1 and the AT8/PHF1 tau conformations implies that the posttranslational modification code of tau contains redundancy, with multiple possible sets of posttranslational modifications leading to the same structure. We hypothesize that in brain, this redundancy and dominance may be crucial for allowing specific pathological folds to emerge from the heterogeneous environment of the cell.

**Figure 6.**
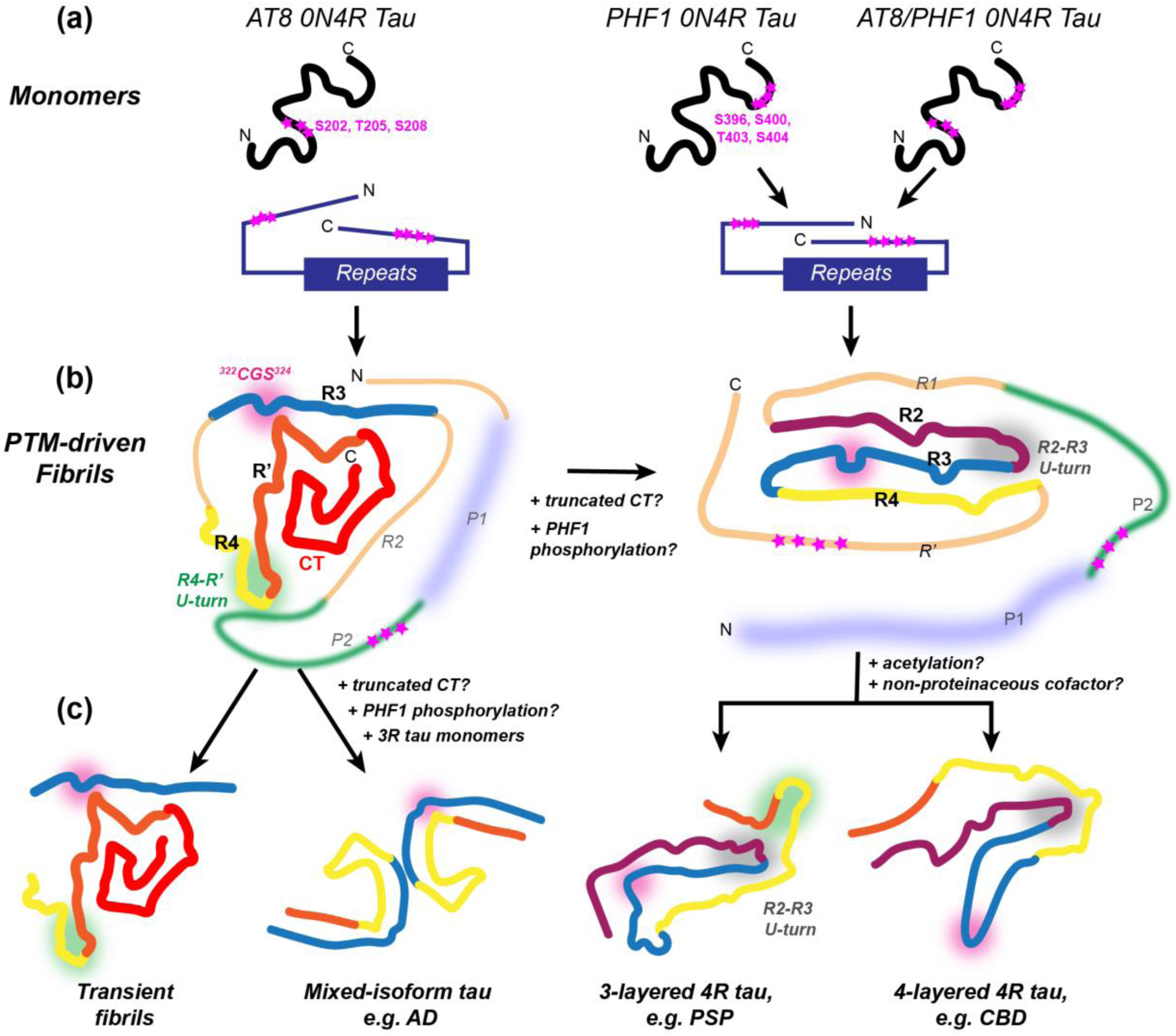
Proposed aggregation pathways of AT8 and PHF1 phosphorylated tau. (**a**) Schematic of soluble tau folds bearing AT8, PHF1, and AT8/PHF1 phospho-mimetic mutations based on FRET data ^44^. (**b**) Schematic of the AT8 tau and AT8/PHF1 tau structures and their fuzzy coat arrangement. Color scheme of the dynamic portions is the same as in Fig. 5d. (**c**) Proposed model of how AT8 tau and PHF1 tau may evolve into brain tau aggregates under various environmental conditions by imperfect templating. The AT8 tau structure may evolve into pathological aggregates such as mixed-isoform AD and CTE tau by the addition of 3R tau, truncated CT monomers, and PHF1 phosphorylated monomers. The AT8 tau structure may also remain as a transient structure in normal brains that can be cleared by phosphatases. The PHF1 tau structure may evolve into brain 4R tau structures such as in PSP and CBD by the addition of acetylated monomers or non-proteinaceous cofactors. Pink shade indicates the flexible ^322^CGS^324^ segment, green shade indicates the R4-R’ U-turn, and grey shade indicates the R2-R3 U-turn.

### Pathological and physiological phosphorylation of tau

The high-resolution structures of these pseudo-phosphorylated tau fibrils provide a structural framework for understanding how tau phosphorylation might lead to pathological tau aggregates, as well as how non-pathological aggregates could exist transiently. Tau is highly phosphorylated not only in tauopathy brains ^23, 25, 30, 53^ but also in normal brains in a developmentally controlled manner ^34, 54–55^. Differentiating pathological and physiological tau phosphorylation requires detailed knowledge of the distribution of phosphorylated residues, extent of phosphorylation at each residue, the sequence of phosphorylation events, and the spatial distribution of phosphorylated tau, in diseased and normal brains. Fetal human tau is highly phosphorylated but with subtle differences from AD tau ^34^. For example, antibody staining data show that S202, S396 and S404 are phosphorylated in both fetal and AD brains whereas pT205 and pS208 are more prevalent in AD tau ^35, 56^. Mass spectrometry data show that pT205 but not the other AT8 residues is enriched in seeding-competent fractions of AD tau ^57^, suggesting that the phosphorylation pattern in the Pro-rich region may differ in specific ways between diseased and normal brains. Tau is also phosphorylated in hibernating animals at some of the AT8 and PHF1 residues such as S199, S202, and S404, but the phosphorylation is reversible after animal arousal ^36, 58^, and the phosphorylated sites correlate with those enriched in sarkosyl-soluble and seeding-incompetent fractions of AD tau ^57^, consistent with the fact that these phosphorylation events do not have deleterious consequences. These data on physiological tau phosphorylation are consistent with emerging mass spectrometry data of the order of phosphorylation during the progression of AD ^57^. The latter data suggest that phosphorylation of N-terminal residues such as S198, S199 and S202 occurs early and exists in sarkosyl-soluble fractions of AD, whereas phosphorylation of S396, S400, T403 and S404 in the C-terminal region occurs in advanced stages of AD. In addition, AT8-positive tissues are preferentially enriched in pretangles whereas PHF1-positive inclusions are preferentially enriched in intracellular and extracellular neurofibrillary tangles ^59^. AT8 immunoreactive neurons have been observed from 1 to 100 years, increasing with age ^4^. Taken together, these biochemical evidence suggests that phosphorylation near the AT8 and PHF1 epitopes are involved in both pathological aggregates and physiological tau. This complexity should be taken into account in understanding the pseudo-phosphorylated tau structures obtained here.

### Comparison of phospho-mimetic tau with brain tau and proposed model of phosphorylated tau aggregation in human brains

The AT8 tau structure exhibits similarities with tau structures in Pick’s disease, globular glial tauopathy (GGT), and CBD-seeded tau in SH-SY5Y cells ^60^ (**Fig. 6c, Fig. S6c**). Residues ^360^Ile-His^374^ forms a U-turn that match closely among these folds, suggesting a common steric zipper between the R4 and R’. The AT8 tau structure is also interesting because as a 4R tau, its rigid core excludes the R2 domain. This absence of R2 resembles mixed-isoform AD and CTE tau structures, which exclude the R2 domain because the 3R tau monomers in these fibrils lack the R2 domain ^61^. Based on these considerations, we propose that the AT8 tau fold could evolve into a mixed-isoform AD-like structure when 3R monomers are incorporated into the growing fibril, and when PHF1 phosphorylated monomers or CT-truncated monomers are added onto the seed, disrupting the CT from the triangular fold (**Fig. 6c**). Under these conditions, the R3 domain would be forced to pack against other segments of the protein such as R4 and R’, leading to an AD-like conformation. Alternatively, under physiological conditions, the AT8 tau structure can remain transiently, without deleterious consequences, and may be cleared by the cell after dephosphorylation by phosphatases, which are highly active in the developing brain and in hibernating animals after arousal.

A similar process of imperfect templating may evolve the three-layered PHF1 tau structure into pathological brain 4R tau aggregates (**Fig. 6c**). The PHF1 tau structure already displays substantial similarities with various brain 4R tau structures ^13^. The R2-R3 U-turn in PHF1 tau is present in CBD and GGT tau, and the ^301^PGGG^304^ turn that separates the R3 and R4 repeats in PHF1 tau is also found in the three-layered PSP tau. In PHF1 tau as well as AT8 tau structures, the ^322^CGS^324^ motif forms part of an inverted Ω, which leaves R3 straight as a whole. In comparison, in three-layered PSP, GGT, and GPT tau aggregates, the ^321^KCGS^324^ segment forms a 90° turn, causing a spacious region that has been hypothesized to contain non-proteinaceous cofactors. The addition of such cofactors could be one mechanism for changing the PHF1 tau fold into a diseased tau structure. If, however, the non-proteinaceous density is lysine acetylation, then we can imagine that the PHF1 tau structure may develop into pathological folds by templating with monomers that contain acetylated lysines. This imperfect templating has been reported for tau aggregates obtained from SH-SY5Y cells seeded with AD PHF tau and CBD brain extracts, and structural differences between the aggregates in the seed and the cell were found ^60^.

### Shape and dynamics of the fuzzy coat in AT8 and PHF1 tau fibrils

In addition to the rigid core structures, NMR spectra that selectively detect the highly mobile residues, together with previous fluorescence resonance energy transfer (FRET) data ^44–45^, yielded information about the dynamics and shape of the fuzzy coat surrounding the rigid cores. Unmodified monomeric tau is known to adopt a paperclip conformation in which the CT lies in the center, flanked by the NT on one side and the MT-binding repeats on the other ^62^. FRET data showed that installation of Glu mutations at S199, S202 and T205 swung the NT outward, separating it from the CT and loosening the paperclip within the monomeric state of tau (**Fig. 6a**). Installing two Glu mutations at S396 and S404 caused the CT to move closer to the NT. Installing all five Glu mutations caused the NT to approach the microtubule-binding repeats as well as the CT, tightening the paperclip. Since the residual conformation of soluble tau contributes to the nucleation of fibrils, and the chemical shifts of the fuzzy coat residues between fibrillar tau and soluble tau are in good agreement ^38, 41–42, 61^, we can approximate the dynamic conformations of the fuzzy coat in the fibrils by those of soluble tau (**Fig. 5d**). In AT8 tau fibrils, we hypothesize that the partially immobilized NT stacks against R3, whereas the partially immobilized R2 shields the CT. The P1-P2 junction lie further on the fibril exterior. In the PHF1 tau fibrils, the semi-mobile NT may stack against R’, which in turn lies outside R4. In the joint mutant, the negative charges at the AT8 epitope may repel the NT from the fibril core, thus explaining the increased dynamics of the NT observed in the INEPT spectra.

In summary, we have determined the structures of two pseudo-phosphorylated full-length tau fibrils by solid-state NMR and cryo-EM. We propose that these PTM tau structures may evolve under cellular and molecular selection pressures ^63^ such as isoform composition and monomers with additional PTMs into disease-relevant structures. They may also exist as transient structures that are reversible in normal brains. Our results suggest that the distinct tau conformations in tauopathies may be caused by distinct patterns of PTMs ^64^. Because brain tau aggregates have variable isoform compositions and heterogeneous PTMs due to the incorporation of differently modified monomers and due to PTMs added post-assembly, the structures of homogeneously pseudo-phosphorylated tau fibrils obtained here are important for understanding the nucleating structures that could evolve into pathological aggregates by imperfect templating.

The integration of solid-state NMR and cryo-EM in this study is synergistic for understanding both the structures of the rigid core and the dynamics of the fuzzy coat. The remarkable agreement between the chemical-shift determined β-strand locations in the amino acid sequence and the cryo-EM densities indicate that each phospho-mimetic tau sample adopts a single predominant molecular structure under our experimental conditions. These structures are readily determined by helical reconstruction, whereas information about the water accessibility of the rigid core and the dynamics of the fuzzy coat is readily obtained from solid-state NMR.

## Material and Methods

### Cloning, Protein Expression and Protein Purification

The genes for AT8-0N4R tau, PHF1-0N4R tau, and AT8/PHF1-0N4R tau were formed by modifying a 0N4R tau gene (Genescript) in a pET-28a vector with Gibson Assembly site-directed mutagenesis. The final expressed protein is 0N4R tau (Uniprot P10636-6) with mutations S202E, T205E, S208E for AT8-0N4R tau, and mutations S396E, S400E, T403E, S404E for PHF1-0N4R tau, and all seven sites for the combined AT8-0N4R, PHF1-0N4R tau. Amino acid numbering follows the 2N4R tau sequence (Uniprot P10636-8), so K44 is followed by A103. Success of mutagenesis was confirmed by Sanger sequencing before the modified plasmids were transfected into *E. coli* BL21(DE3) competent cells. Colonies from freshly transformed LB Agar plates were used to inoculate a 10 mL LB starter culture. After overnight growth at 37°C with 250 rpm shaking, the starter culture was used to inoculate 1 L LB media, which was allowed to grow at 37°C with 250 rpm shaking to an OD_600_ of 0.8. At this point the cells were spun down at 1,000 *g* for 20 minutes. The cell pellet was resuspended in 1 L M9 minimal media containing 2 g/L ^13^C_6_-D-glucose and 1 g/L ^15^NH_4_Cl as the only carbon and nitrogen sources, respectively. After 30-60 minutes of shaking at 250 rpm and 37° C, the cells reached an OD_600_ of 1.0, at which point protein expression was induced with 1 mM IPTG, and an additional 1g/L ^13^C_6_-D-glucose was added to the media. After 3-5 hours of protein expression at 37°C, cells were harvested by centrifugation and pellets were frozen. The LB Agar plates and all media contained 50 μg/mL Kanamycin.

The phospho-mimetic tau proteins were purified using similar procedures as for wild-type tau ^41–42, 65^. Briefly, cells were thawed and homogenized by vortexing in ice-cold lysis buffer, which contains 20 mM Na_2_HPO_4_ (pH 6.8), 50 mM NaCl, 5 mM DTT, 0.1 mg/mL lysozyme, and 1x cOmplete™ protease inhibitor cocktail tablet (Roche) per 50 ml lysis buffer. After lysis on ice with a probe sonicator, 5 s on/5 s off for 10 mins, the lysate was placed in a boiling water bath for 20 minutes, then centrifuged at 15,000 *g* for 40 minutes to remove cell debris and aggregated protein. The supernatant was purified with a cation-exchange column (self-packed with SP Sepharose Fast Flow resin, GE healthcare) running between 20 mM Na_2_HPO_4_ (pH 6.8), 50 mM NaCl, 2 mM DTT, and a gradient with the same buffer but with 1 M NaCl. Eluent fractions containing tau were further purified by reverse-phase HPLC (Agilent Zorbax 300SB-C3 column, 21.2 x 250 mm, 7 μm particle size) using an acetonitrile gradient of 5-50% in 50 min. HPLC fractions were pooled and lyophilized. Typical yields were ∼40 mg purified tau per 1 L M9 medium for all three mutants.

### Cofactor-Free Fibrillization

Lyophilized PHF1-0N4R tau was dissolved at 1.6 mg/mL in 50 mM K_2_HPO_4_: KH_2_PO_4_ buffer, pH 6.8, containing 300 mM NaCl, 5 mM DTT, and 1x cOmplete™ protease inhibitor cocktail tablet (Roche) per 40 ml fibrillization buffer. AT8-0N4R tau was fibrillized identically but at a protein concentration of 0.4 mg/mL. AT8/PHF1 tau was fibrillized identically but at a protein concentration of 1.6 mg/mL and 150 mM NaCl.

Freshly dissolved tau was sonicated for 10 minutes to ensure complete dissolution, then the solution was transferred to a 500 mL glass bottle containing 15-30 mL solution, which had been flushed with nitrogen gas to remove oxidizing O_2_. The solution was shaken at 250 rpm (orbital diameter 25 mm) at 37°C for 14 days. To maintain a reducing environment for the two cysteine residues in the protein (C291 and C322), 2 mM DTT was added every 2 days, with N_2_ flushing each time the reaction vessel was opened. After 14 days, aliquots of the fibrillization reaction were taken and frozen to be used for TEM and cryo-EM experiments, while the remaining solution was pelleted at 100,000 *g* for 1 hour at 4°C in a TLA-55 rotor. The pellet was packed into a 3.2 mm pencil-style MAS NMR rotor using centrifugation and manual packing with a needle. Additional water was centrifuged into the packed rotor as needed to ensure uniform hydration.

### Negative-Stain Transmission Electron Microscopy

Unconcentrated tau fibril solutions were adsorbed onto freshly glow-discharged, 200-mesh formvar/carbon-coated copper grids (Ted Pella), extensively washed with water to remove any trace of phosphate buffer, and stained with 0.7% (w/vol) uranyl formate for 15-30 s. TEM images were taken on an FEI Tecnai T12 electron microscope.

### Solid-State NMR Spectroscopy

Solid-state NMR experiments were conducted on a Bruker Avance II 800 MHz (18.8 T) spectrometer in the Francis Bitter Magnet Lab (FBML) at MIT, with a BlackFox 3.2 mm HCN MAS probe. ^13^C chemical shifts were referenced externally to the adamantane CH_2_ chemical shift at 38.48 ppm on the tetramethylsilane (TMS) scale, and ^15^N chemical shifts were referenced to the ^15^N peak of ^15^N-acetylvaline at 122.0 ppm on the liquid ammonia scale. Typical radiofrequency (rf) field strengths were 50-83 kHz for ^1^H, 50–62.5 kHz for ^13^C, and 25–36 kHz for ^15^N. ^1^H chemical shifts of a hydrated DSS-containing POPC membrane sample was used as an external standard to calibrate the ^1^H chemical shifts and measure the sample temperature. Reported temperatures are estimated sample temperatures based on the probe thermocouple and the measured water ^1^H chemical shift ^66^. Experiments were conducted under 10.5 kHz or 14 kHz MAS (**Table S4**).

Rigid residues were detected using dipolar polarization transfer experiments such as ^13^C-^13^C CORD and ^15^N-^13^C SPECIFIC-CP. 2D ^13^C-^13^C spin diffusion correlation spectra were measured using ^1^H-^13^C cross polarization (CP) followed by ^13^C-^13^C CORD mixing ^67^. Additionally, DREAM ^68^, or ^13^C-^13^C cross polarization, was used for ^13^C polarization transfer in some of the 2D CC spectra. 2D NCACB correlation spectra were measured using ^1^H-^15^N CP followed by ^15^N-^13^Cα ^SPECIFIC^CP ^69^ and ^13^C-^13^C DREAM transfer ^68^. DREAM transfer was accomplished by a 50-100% amplitude ramp from 4 kHz to 8 kHz for ^13^C spin lock at a carrier frequency of 55 ppm. This large ramp amplitude allowed the Cα magnetization to be efficiently transferred to the serine Cβ peaks at ∼62 ppm, an unusually low alanine Cβ peak at 12 ppm, in addition to the Cβ sites at the common chemical shifts between these two extremes.

Three-dimensional dipolar correlation spectra were measured to assign the resonances or rigid residues in AT8-0N4R tau and PHF1-0N4R tau (**Table S4**). For PHF1-0N4R tau, we measured 3D NCACX, NCOCX and CONCA correlation spectra. The NCACX and NCOCX spectra used 4-5 ms ^15^N-^13^C ^SPECIFIC^CP transfers and 82 ms for ^13^C-^13^C CORD spin diffusion. The CONCA experiment used two and ^SPECIFIC^CP transfer steps from CO to ^15^N and from ^15^N to the bonded CA. For AT8-0N4R tau, which has a larger rigid core, in addition to these three 3D spectra, we also measured a 3D CAN(CO)CA spectrum and a 3D CONCACB spectrum. The CAN(CO)CA experiment used a modified ^BSH^CP sequence ^70 41^ for CO-Cα magnetization transfer while the CONCACB experiment used the DREAM sequence for Cα-Cβ magnetization transfer. The latter provided many useful correlation peaks to disambiguate resonance assignment.

For many 2D and 3D experiments, a short ^1^H-^13^C CP contact time of 70 μs was used to select only the most rigid β-sheet signals while excluding the more dynamic signals from partially mobile regions of the protein. Experiments requiring significant signal averaging to achieve sufficient signal-to-noise ratios (SNR) were run in blocks of 1-3 days with field drift correction between each block. Multiple blocks of repeated 2D or 3D spectra were added in the time domain before Fourier transformation.

The 2D and 3D MAS NMR spectra were acquired and processed in TopSpin 3.2, and chemical shift assignment was conducted in SPARKY. Typical processing in TopSpin used either QSINE apodization with SSB = 3 or GM apodization with LB = –20 Hz and GB = 0.05. Spectra were plotted with 1.2 x multiplication factor between successive contour lines.

Water-edited 2D ^13^C-^13^C correlation spectra were measured at 14 kHz MAS using a water-selective ^1^H spin echo consisting of a 1 ms 180° Gaussian pulse surrounded by one rotor period before and after the pulse. This water-selective T_2_ filter is followed by a ^1^H mixing time of 4 ms to allow water ^1^H magnetization to transfer to hydrated protein ^1^H, after which standard 2D CC or NCACB experiments are conducted. Without the ^1^H spin diffusion period, less than 0.1 % of the protein signal is observed, indicating that the ^1^H T_2_ filter removed all rigid protein ^1^H magnetization. With a 100 ms ^1^H spin diffusion period, 1D spectra matched the unedited spectra, except with 50% of the total intensity for AT8-0N4R tau or 55% of the total intensity for PHF1-0N4R tau. Since the 100 ms water-edited spectra and the unedited spectra matched except for an overall intensity difference, 2D control spectra were acquired without the T_2_ filter, and were instead scaled to 50% or 55% intensity for AT8-0N4R or PHF1-0N4R tau, respectively. For resolved peaks in the 2D spectra, the intensities of the 4 ms water-edited spectrum (S) and the unedited spectrum (S_0_) were extracted, summed with intensities of other resolved peaks for the same residue, divided (S/S_0_) and then corrected for the number of scan difference. Error bars were propagated from the SNRs of 1D cross sections of resolved peaks in the 2D spectra. For PHF1-0N4R tau, water-edited NCACB spectra were acquired with the same water-selective ^1^H filter as the 2D CC spectra and analyzed identically. When both NCACB and CC DREAM spectra gave the S/S_0_ values for the same residue, the two ratios were averaged so to minimize the propagated error of the average S/S_0_ value.

Mobile residues were detected using J-based polarization transfer experiments, including ^1^H-^15^N refocused INEPT, ^1^H-^13^C refocused INEPT, and ^1^H-^13^C INEPT followed by ^13^C-^13^C TOCSY. TOCSY was conducted with 12 ms of 36 kHz DIPSI-3 on the ^13^C channel, with the rf carrier frequency set to 50 ppm. No ^1^H decoupling was necessary during TOCSY transfers.

### Chemical shift assignment

The 2D and 3D dipolar correlation spectra for the proteins were sequential assigned following standard protocols. 2D CC, 2D NCα, and 3D NCACX spectra were first analyzed to identify the residue types of all rigid and semi-rigid residues. Then the Cα and CO chemical shifts were used in conjunction with 3D NCOCX and CONCA spectra for sequential assignment. A composite metric of the secondary chemical shift difference between Cβ and Cα, Δδ_*C*β_ − Δδ_*C*α_ ≡ (δ_*C*β,*expt*_ − δ_*C*β,*coil*_) − (δ_*C*α,*expt*_ −δ_*C*α,*coil*_), was used to identify the β-strand segments. Positive Δδ_*C*β_ − Δδ_*C*α_ values denote β-strand segments while negative values indicate α-helical segments.

### Cryo-EM data acquisition

3 μl aliquots of the fibrillization reactions were thawed then applied to glow-discharged R1.2/1.3, 300 mesh carbon Au grids for AT8-0N4R tau, and R1.2/1.3, 400 mesh carbon Au grids for PHF1-0N4R tau (SPI Supplies). The grids were plunge-frozen in liquid ethane using a Thermo Fisher Scientific Vitrobot Mark IV system. Cryo-EM data were acquired at the MIT.nano facility. All images were recorded at a dose of ∼50 electrons per Å^2^ using the EPU software (Thermo Fisher Scientific), and converted to tiff format using relion_convert_to_tiff prior to processing. Images were recorded on a Krios G3i with Bioquantum K3 camera (Gatan), using an energy slit of 20 eV on a Gatan energy filter (**Table 1**).

### Helical reconstruction

Helical reconstructions were performed in RELION-4.0^71–72^. The frames of the raw EM movies were adjusted for gain, aligned, weighted, and combined using RELION’s motion correction program^73^. CTFFIND-4.1^74^ was used to estimate the parameters of the Contrast Transfer Function (CTF). Filaments were manually selected in RELION 4.0. The picked particles were initially extracted in boxes of 1028 pixels downscaled to 256 pixels. Reference-free 2D classification was conducted on these particles to assess different variations and crossover distances and to select segments for further processing. The chosen 2D classes were then re-extracted in full-resolution boxes of 512 pixels. Other rounds of reference-free 2D classification were performed using T values of 2-20. 4-20 classes showing clear 4.75 Å cross-β monomer separation were manually chosen and aligned to generate initial models with relion_helix_inimodel2d (**Fig. S5**). Subsequently, several rounds of 3D classification were employed to select particles that led to the best reconstruction, with parameters T=2-20 and K=1-4. 3D auto-refinement was utilized to enhance image resolution and optimize the helical twist and rise parameters. Bayesian polishing^73^ and CTF refinement ^75^ were applied to all rendered maps to further improve the resolution. The final maps were sharpened using standard post-processing techniques in RELION, and the reported resolutions were estimated using a threshold of 0.143 in the FSC between two independently refined half-maps ^76^, utilizing the Phenix software suite (**Fig. S10**). The software Resmap ^77^ was independently used to assess the local resolution of the maps (**Fig. S11**).

### Structure calculation and validation

Atomic structural models were constructed *de novo* using COOT, with three or five rungs for each structure. ISOLDE was used for coordinate refinement ^78–79^. Dihedral angles from the middle rung, which served as a template, were applied to the adjacent rungs above and below. Model refinements were conducted separately for each refined structure by raising the temperature to 300 K for 1 minute. The final model was validated using phenix.comprehensive_validation ^80^. Additional information about data processing, model refinement, and validation can be found in **Table 1**.

## References and Notes

## Author Contributions

N.E.M. and A.J.D expressed, purified, and fibrillized the protein; A.J.D. and N.E.M conducted and analyzed the solid-state NMR experiments; A.J.D., N.E.M., and P.D. conducted cryo-EM experiments and analysis; N.E.M. and A.J.D. wrote the first draft of the paper. M.H. and P.D. edited the paper; all authors analyzed the results and interpreted the results; M.H., N.E.M. and A.J.D. designed the experiments. M.H. supervised the project.

## Data availability

The AT8-0N4R tau structure and cryo-EM densities have been deposited into the Protein Data Bank (PDB) with the entry ID 8TTL and the EMDB with the entry ID 41610. The PHF1-0N4R tau structure and cryo-EM densities have been deposited into the PDB with the entry ID 8TTN and EMDB with the entry ID 41611.

