## Supplementary Tables and Figures for "Structures of AT8 and PHF1 Phospho-Mimetic Tau: Insights Into the Posttranslational Modification Code of Tau Amyloid Formation"

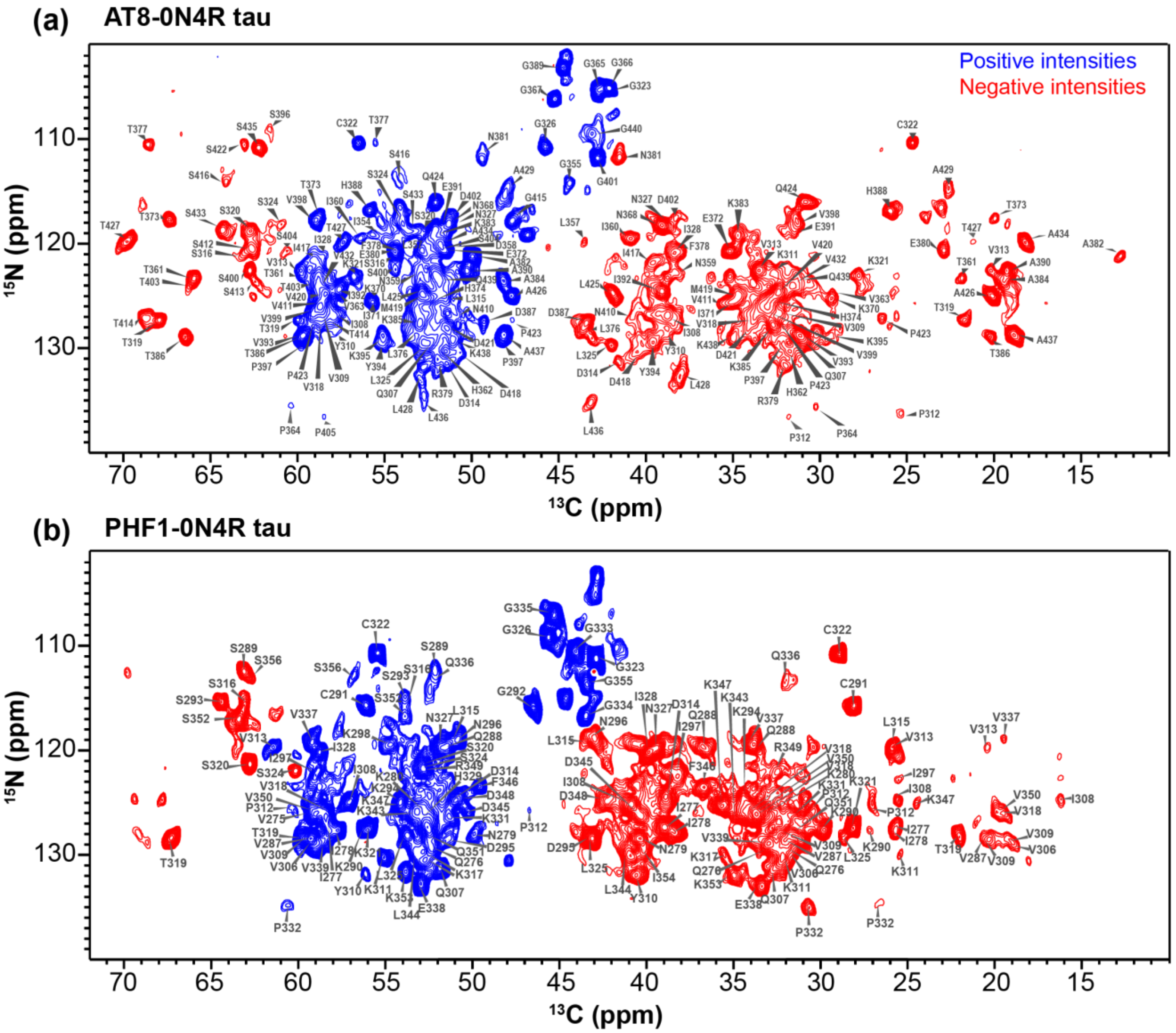


**Figure S1. 2D NCACB spectra of phospho-mimetic full-length 0N4R tau**. (**a**) AT8-0N4R tau. (**b**) PHF1-0N4R tau. Assignments are obtained from 3D correlation spectra. $^{15}$N-$^{13}$C$\alpha$ cross peaks have positive intensities (blue) whereas $^{15}$N-C$\beta$ cross peaks have negative intensities (red) due to the double-quantum DREAM transfer from C$\alpha$ to C$\beta$. A small number of negative $^{15}$N-C$\gamma$ cross peaks are also observed due to direct N-C$\beta$ polarization transfer followed by DREAM C$\beta$-C$\gamma$ transfer.

**(a)** 23 ms CC CORD, AT8-0N4R tau

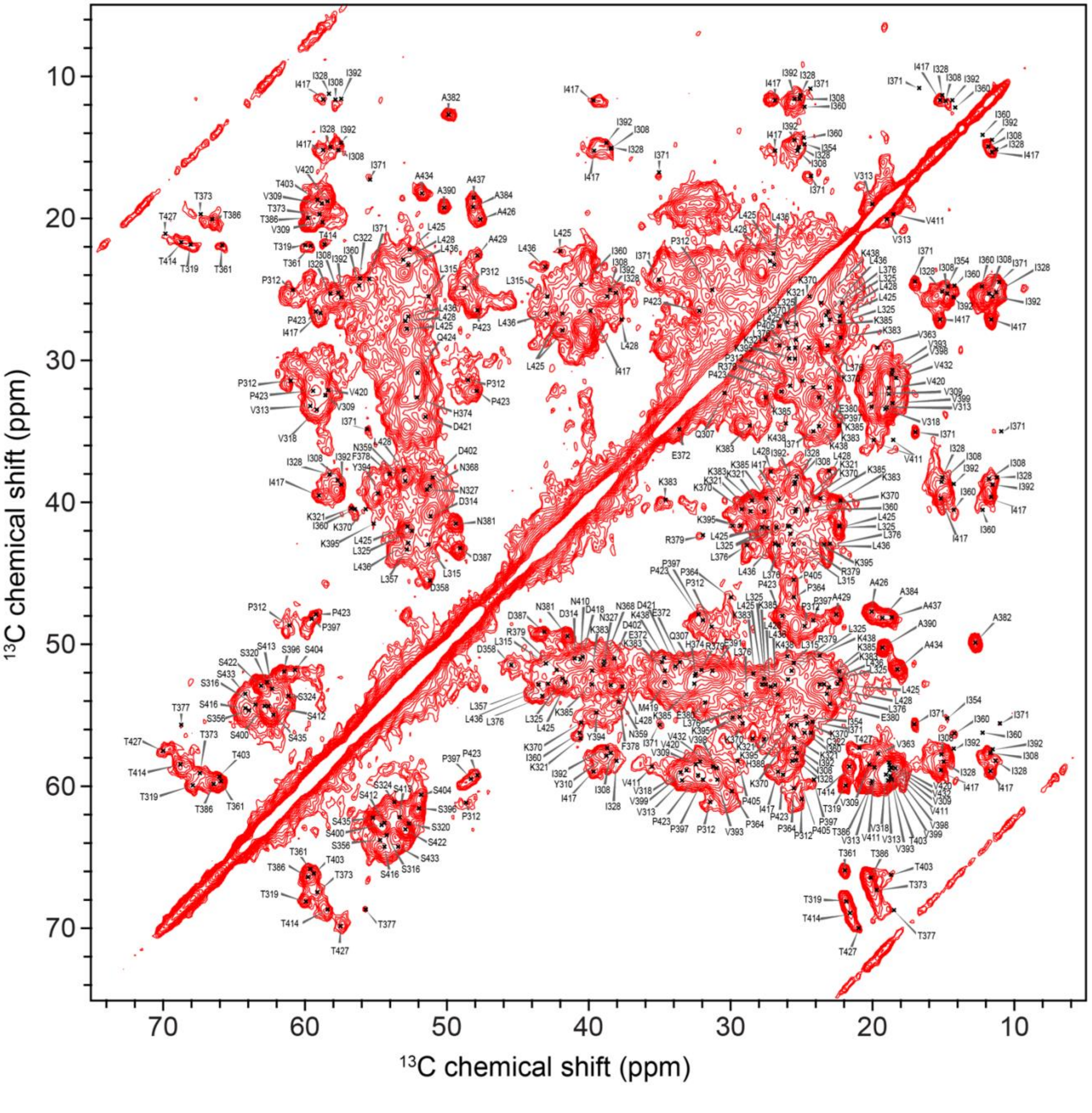

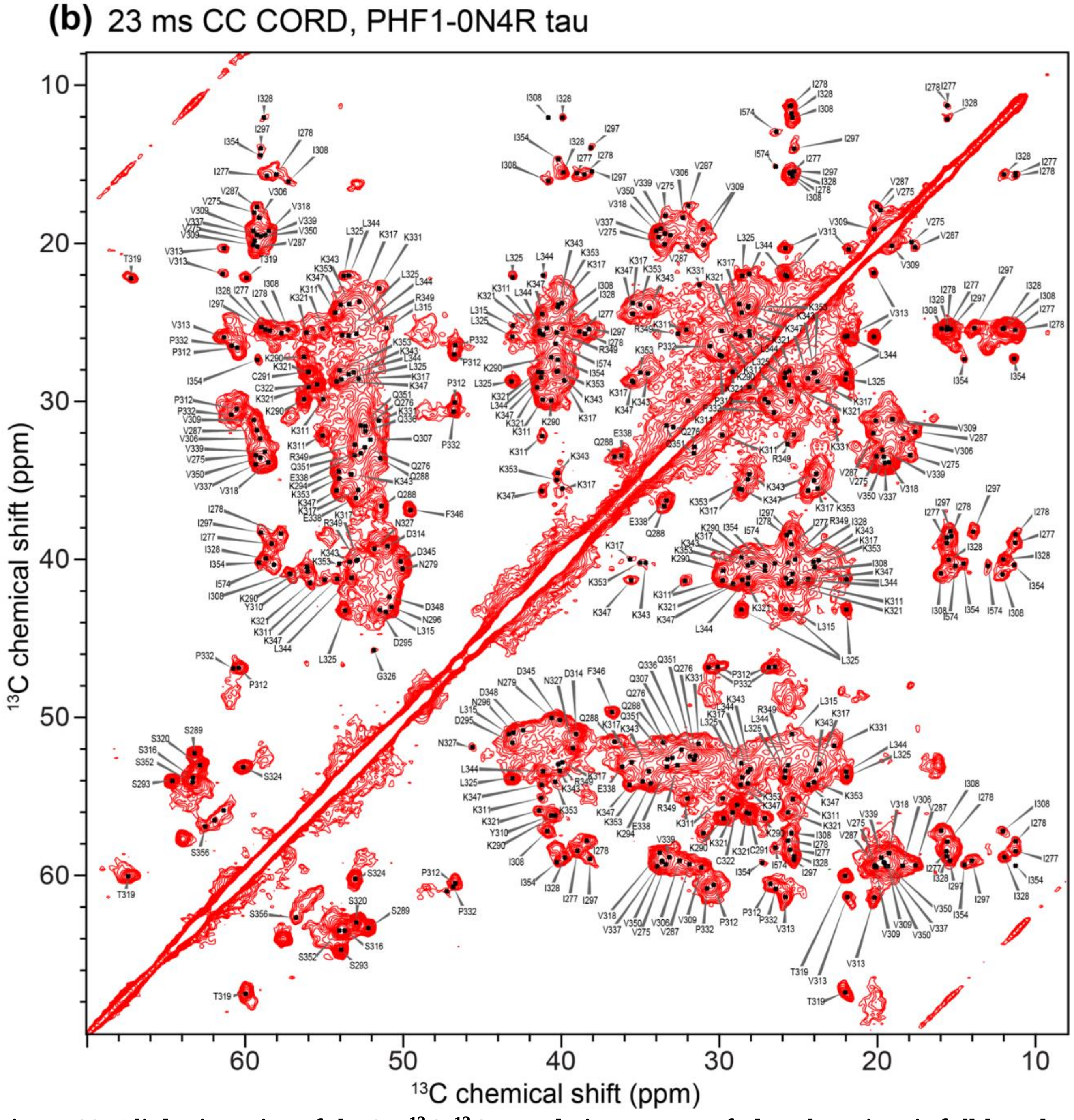


**Figure S2. Aliphatic region of the 2D $^{13}$C-$^{13}$C correlation spectra of phospho-mimetic full-length tau fibrils**. Resonance assignment is obtained from 3D correlation spectra. (**a**) AT8-0N4R tau. (**b**) PHF1-0N4R tau.

(a) AT8-0N4R tau 3D assignment example

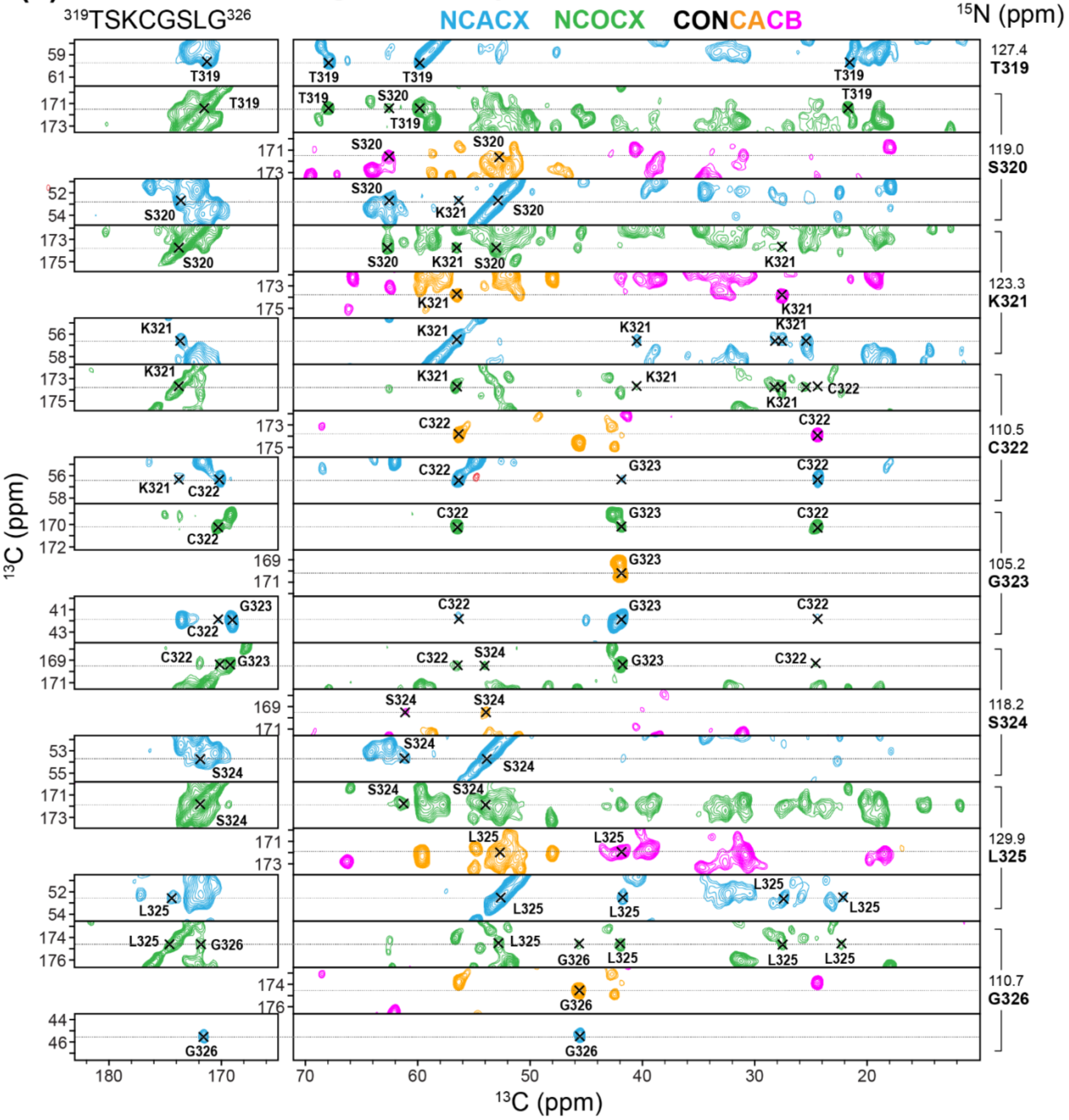

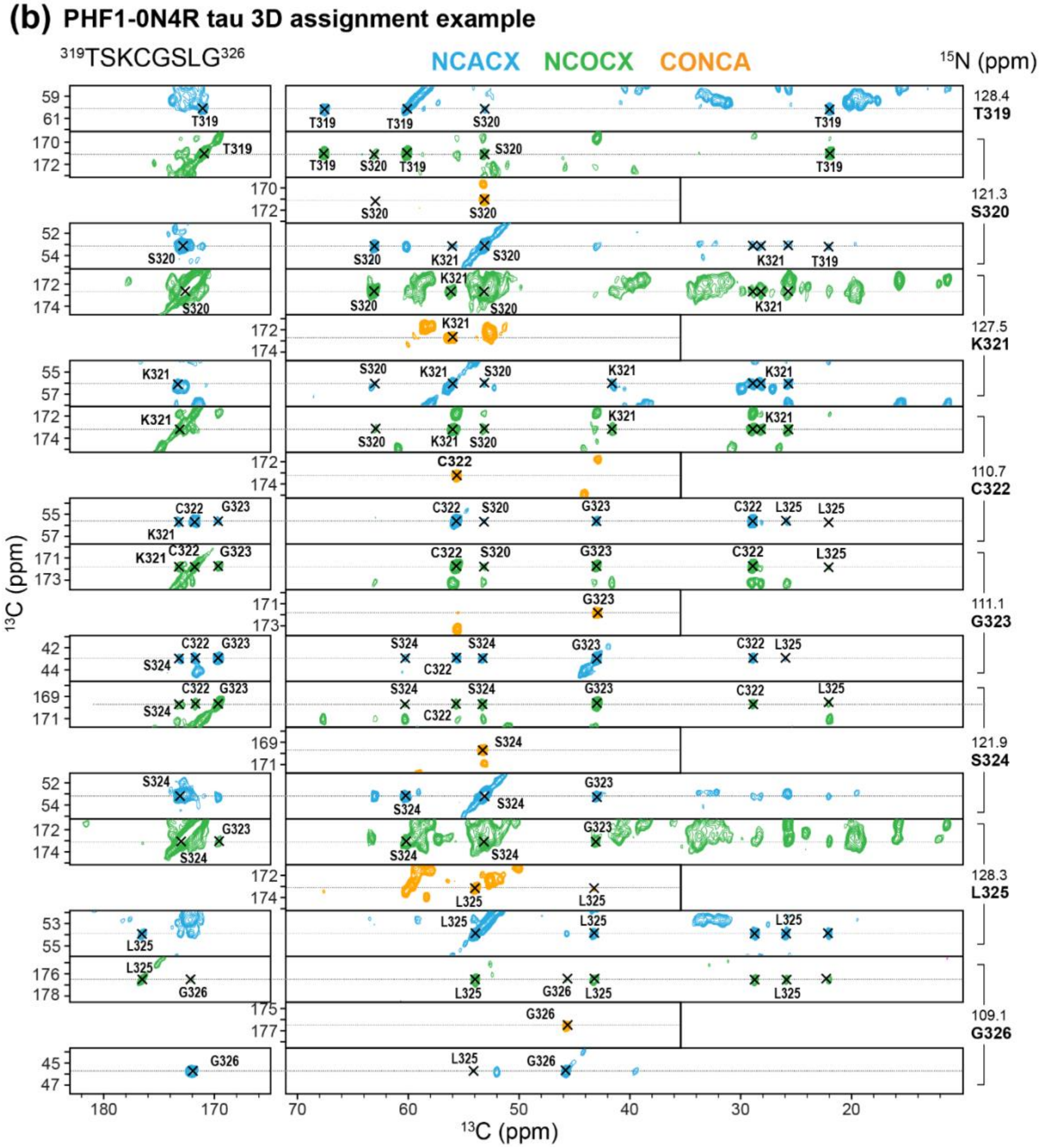


**Figure S3**: **Representative 3D spectral strips of phospho-mimetic 0N4R tau fibrils for $^{13}$C and $^{15}$N resonance assignment**. (**a**) AT8-0N4R tau, showing the assignment of residues T319-G326. (**b**) PHF1 0N4R tau, showing the assignment of the same segment. The NCACX spectrum (cyan) establishes intra-residue correlations, whereas the NCOCX and CONCACB spectra establish sequential correlation peaks.

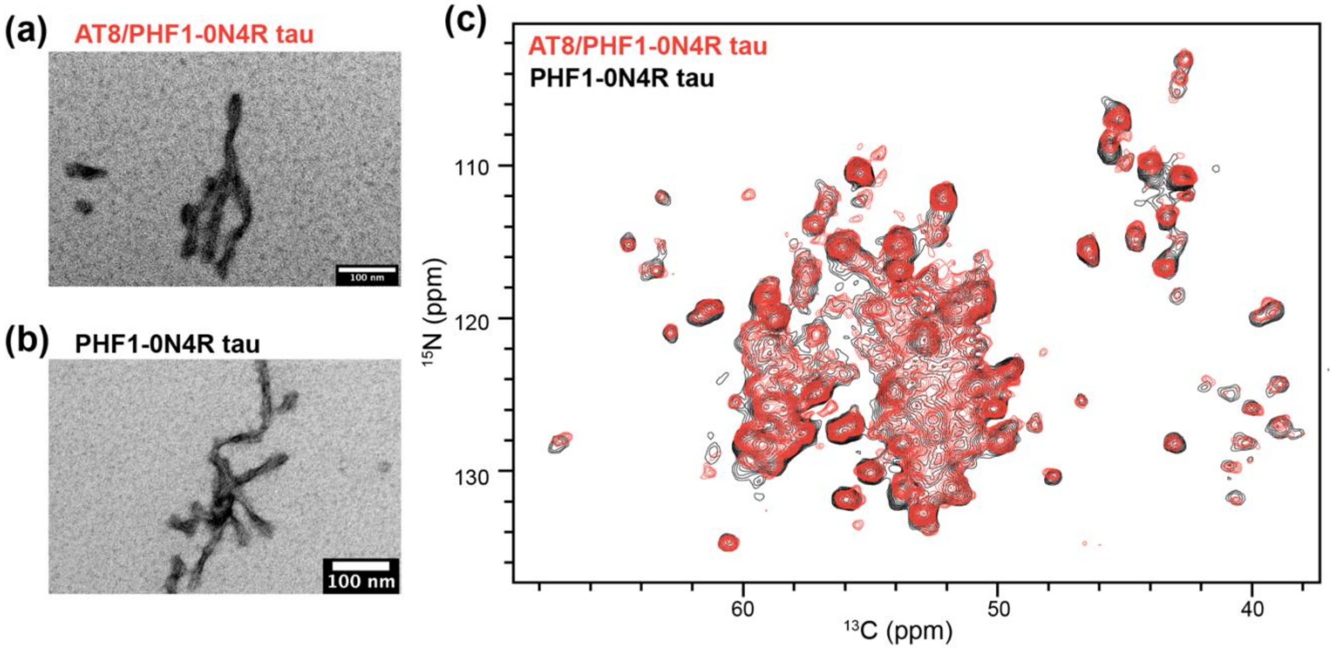


**Figure S4**. **The AT8/PHF1 phospho-mimetic mutant tau has the same fibril core structure as PHF1-only mutant tau**. (**a**) Negative stain TEM images of AT8/PHF1-0N4R tau and PHF1-0N4R tau fibrils. (**b**) 2D NCA spectra of the joint AT8/PHF1 mutant fibril (red) and the PHF1-only mutant. The spectra are nearly identical, indicating that the two samples rigid cores are indistinguishable. Thus, PHF1 pseudo-phosphorylation dominates AT8 pseudo-phosphorylation in dictating amyloid fibril structure.

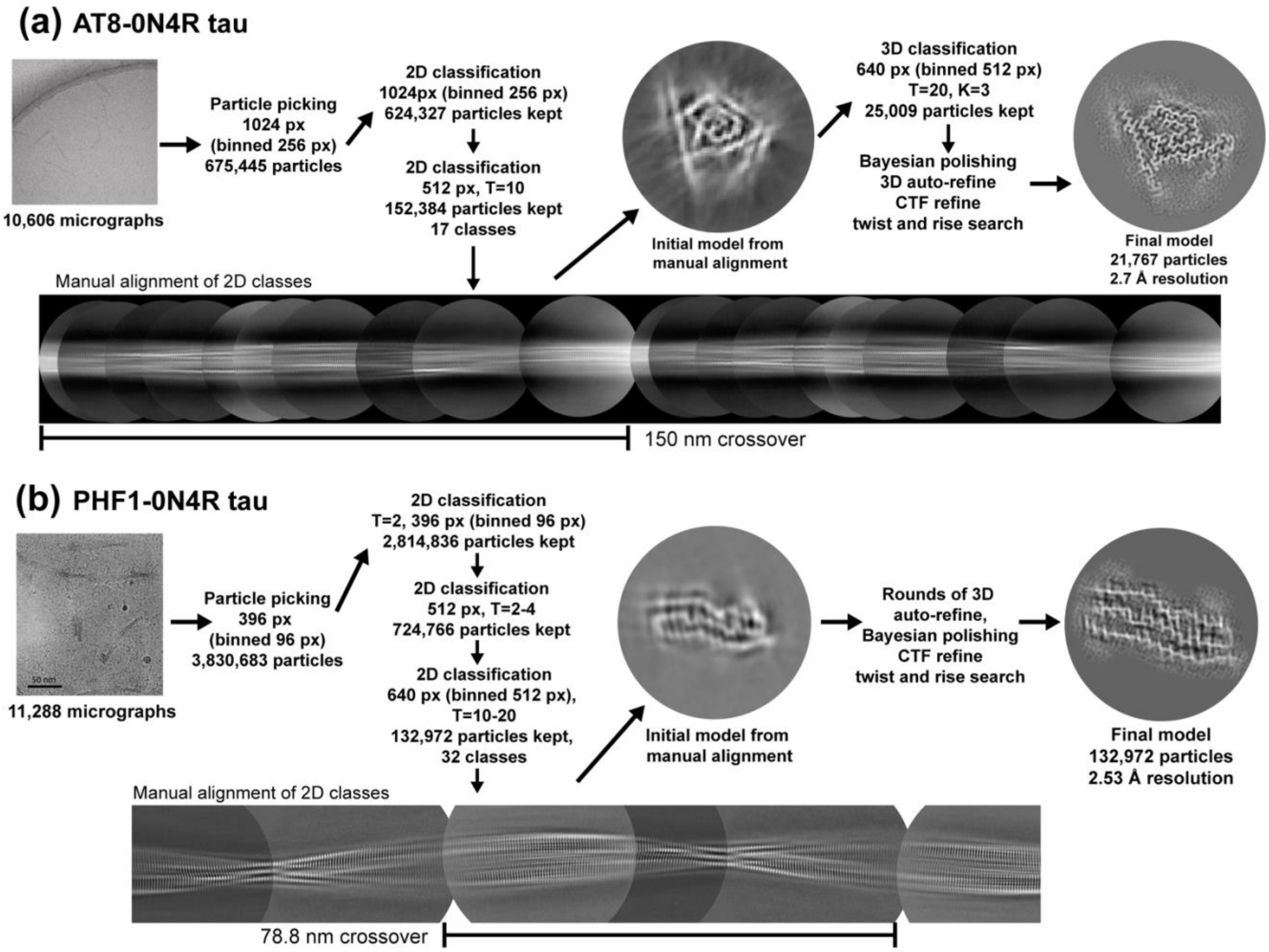


**Figure S5. Cryo-EM workflow for obtain atomic structural models of phospho-mimetic 0N4R tau fibrils**. (**a**) AT8-0N4R tau. (**b**) PHF1-0N4R tau. Both datasets were analyzed through repeated rounds of 2D classification and manual alignment of the 2D classes to obtain accurate crossover lengths.

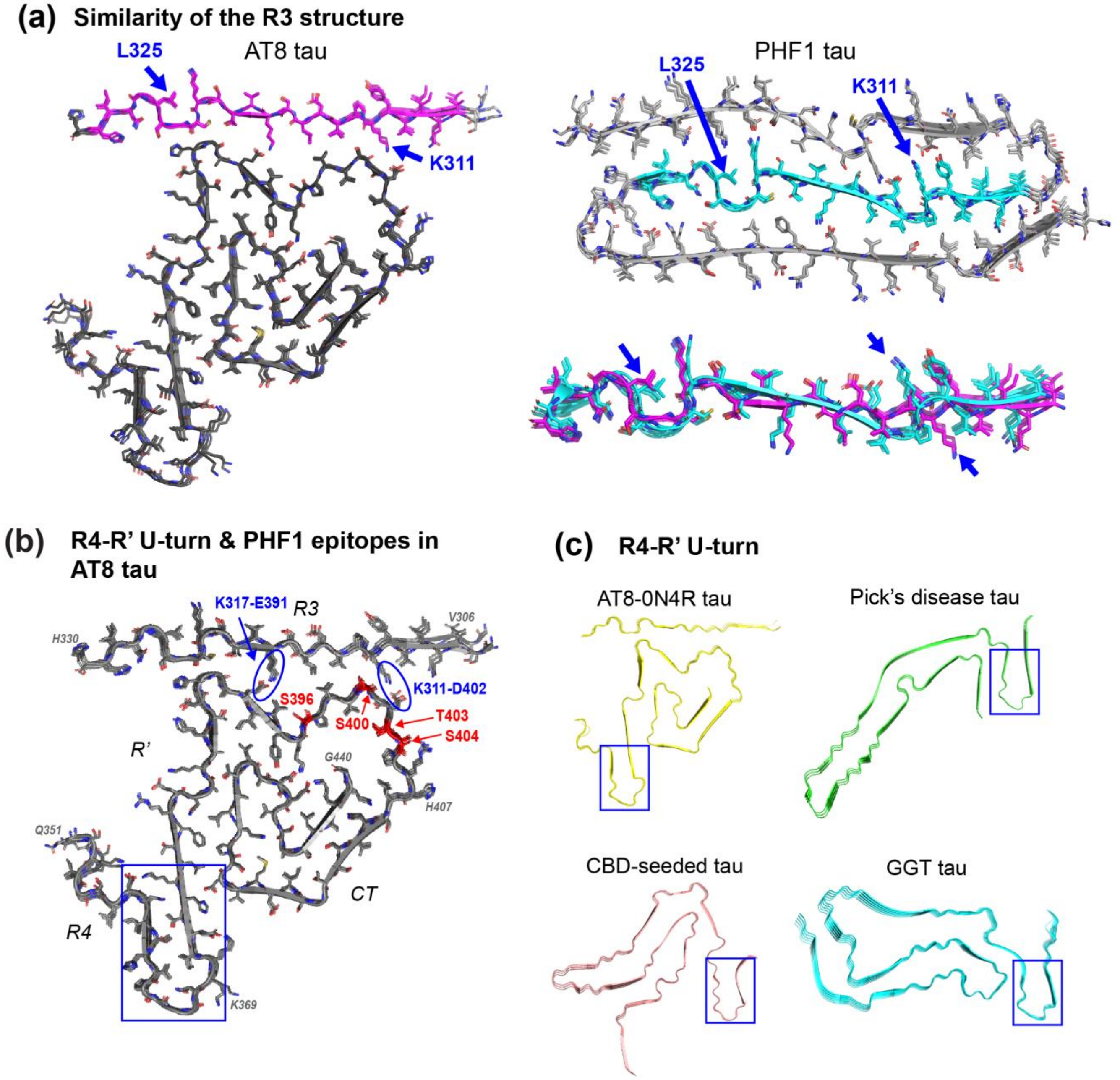


**Figure S6**. **Common structural elements among phospho-mimetic tau and brain extracted tau fibrils**. (**a**) The R3 domain has very similar conformations between AT8- and PHF1-0N4R tau. The only residue with a major structural difference is K311 at the end of the hexapeptide motif, which points to the CT in AT8-0N4R tau but to the R2 in PHF1-0N4R tau. (**b**) Atomic model of AT8-0N4R tau fibril, highlighting the four PHF1 epitope residues that are mutated to Glu in this study. The CT-R3 interaction in AT8 tau is stabilized by K311 and K317 of R3, which form putative salt bridges with E391 and D402, respectively (ovals). Pseudo-phosphorylation of PHF1 residues should disrupt this interaction. The R4-R' U-turn is highlighted by a rectangle. (**c**) The R4-R' U-turn in AT8-0N4R tau is also found in several *ex vivo* tau of both 3R and 4R isoforms.

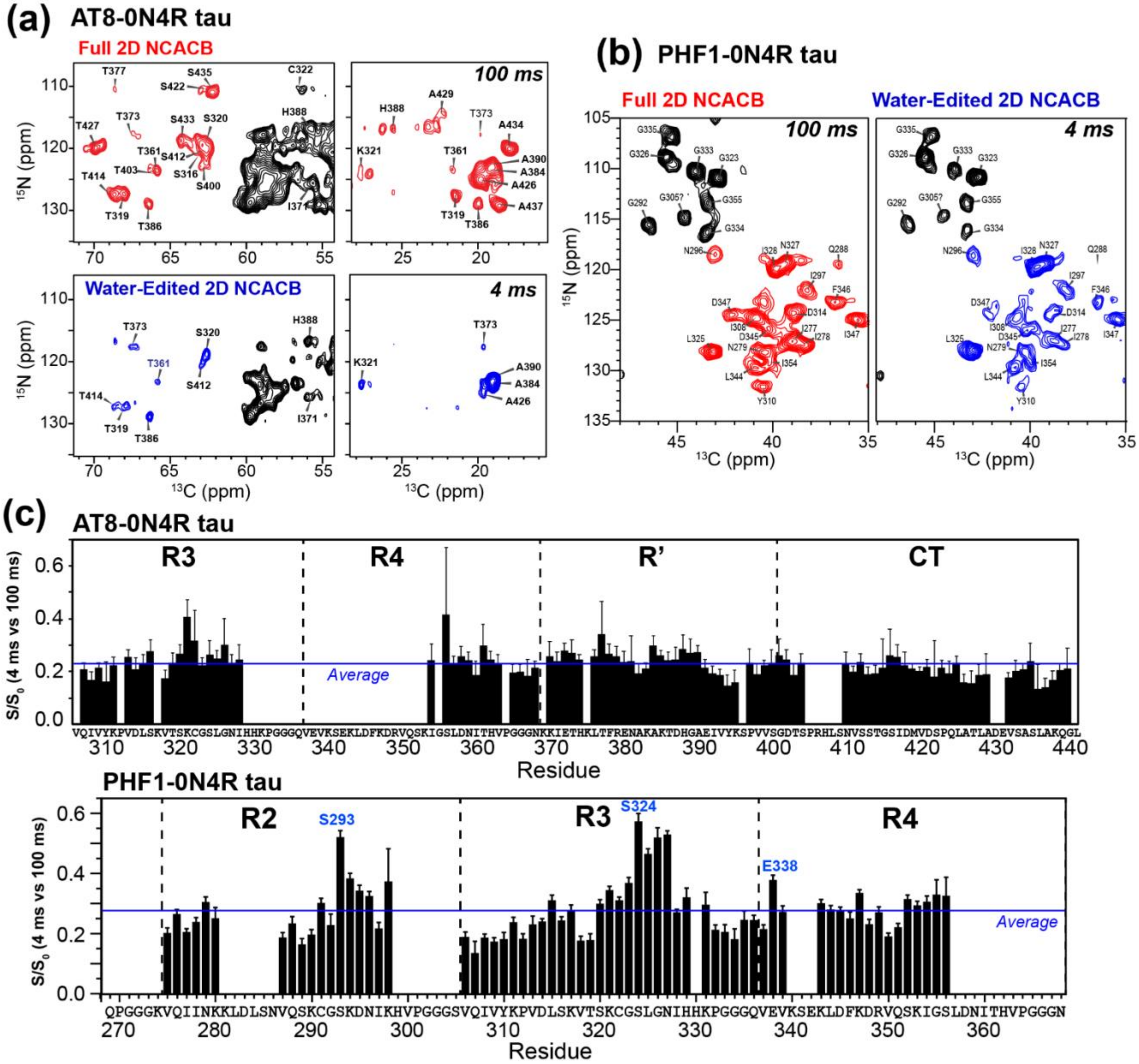


**Figure S7**: Water-edited solid-state NMR spectra for extracting site-specific water accessibilities. (**a**) Representative regions of the water-edited 2D NCACB spectra of AT8-0N4R tau. Positive intensities are shown in black and negative intensities are shown in red (100 ms) or blue (4 ms). (**b**) Representative regions of the water-edited NCACB spectra of PHF1-0N4R tau. (**c**) Water-accessible intensity ratios per residue obtained from a joint analysis of the NCACB and CC-DREAM spectra of PHF1-0N4R tau and the NCACB spectrum of AT8-0N4R tau. Error bars are propagated from the spectral signal-to-noise ratios.

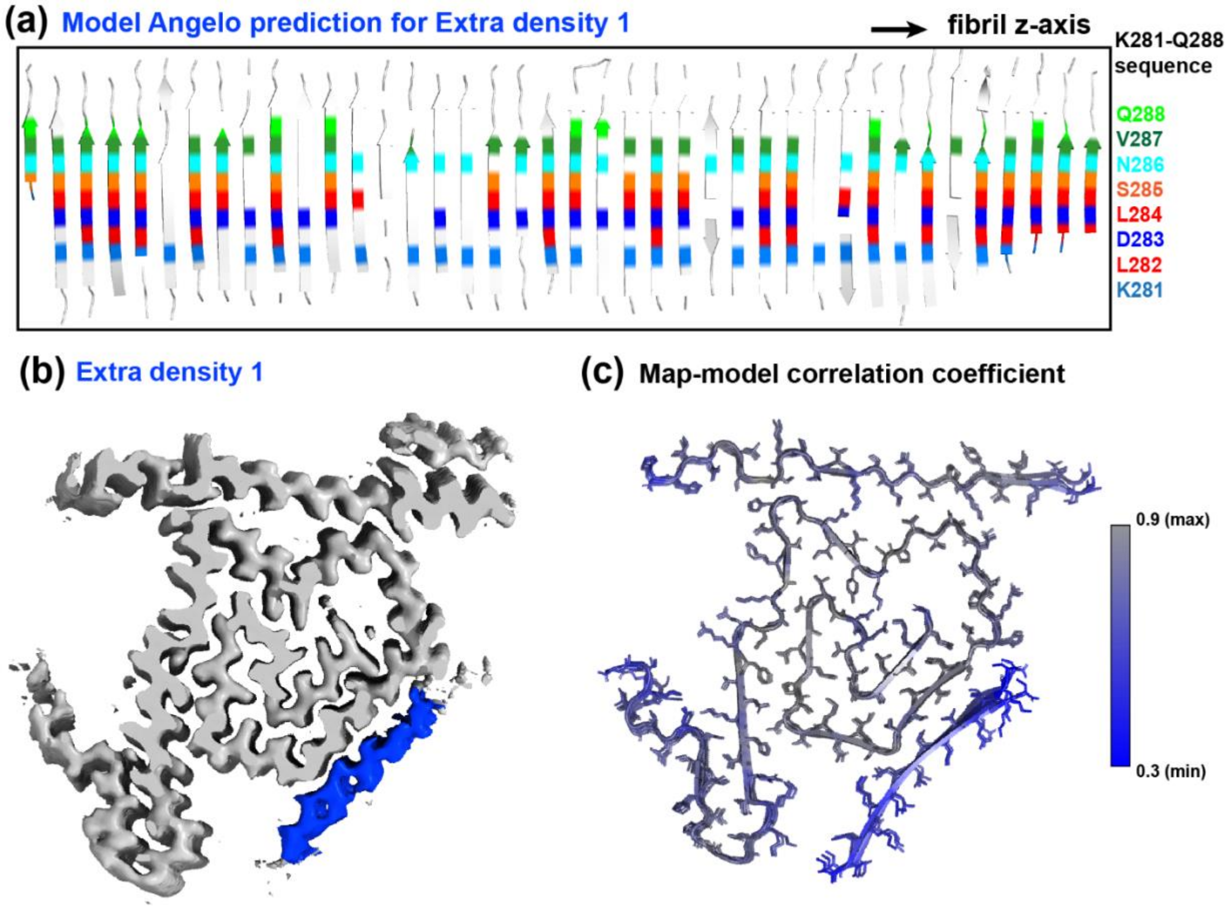


**Figure S8. Tentative assignment of the extra density 1 in the AT8-0N4R tau cryo-EM map using Model Angelo**. (**a**) Prediction of the extra density 1 using about 40 chains along the fibril axis. The consensus prediction is the R2 segment K281-Q288. The predicted residue types are shown in color using the scheme on the right. Residue types that do not correspond to K281-Q288 are shown in white for clarity. (**b**) Extra density 1 (blue) relative to the rigid core of AT8-0N4R tau. (**c**) Correlation coefficients between the atomic model and the refined cryo-EM density map. The predicted K281-Q288 segment has lower correlation coefficients than the rest of the model.

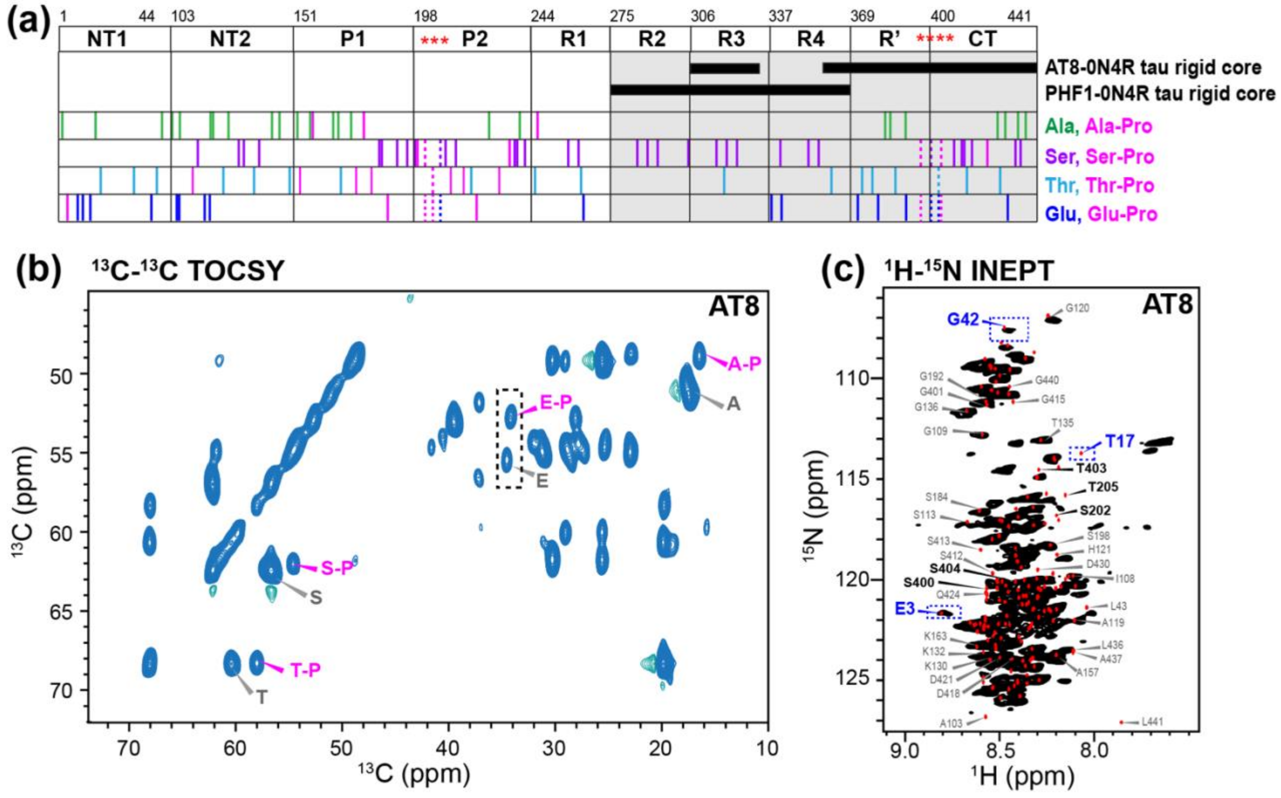


**Figure S9. Fuzzy coat dynamics of pseudo-phosphorylated tau fibrils. (a)** Distribution of Glu, Ser, Thr and Ala residues in 0N4R tau. When a residue precedes a Pro, the residue position is marked by a pink line. (**b**) Region of the 2D $^{13}$C-$^{13}$C TOCSY spectrum of AT8 tau fibrils. (**c**) Region of the 2D $^{15}$N-$^{1}$H INEPT spectrum of AT8 tau fibrils. The regions marked by dashed rectangles are expanded in Figure 4. The other two tau fibrils have qualitatively similar spectral patterns.

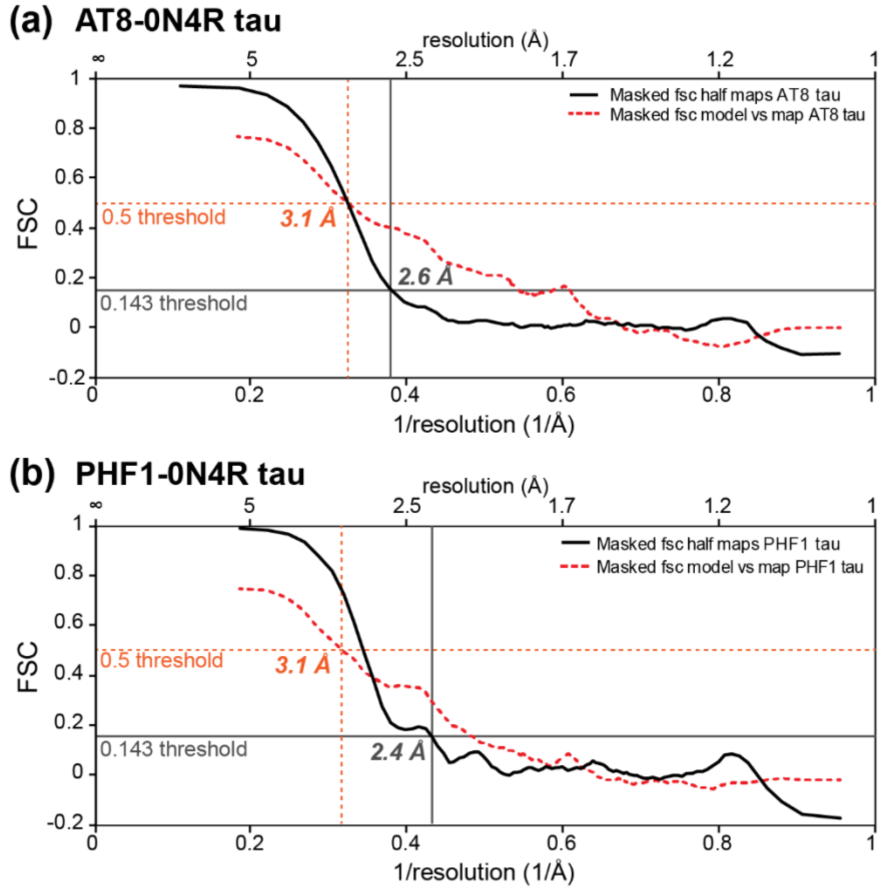


**Figure S10**. **Fourier shell correlation (FSC) of cryo-EM maps and structures to obtain the resolution of the structures**. (**a**) AT8-0N4R tau (**b**) PHF1-0N4R tau fibrils. The FSC curves for two independently refined cryo-EM half-maps are shown in solid line, to which a cutoff value of 0.143 is applied to obtain the resolution. The FSC curves for the final atomic model against the final cryo-EM map are shown in dashed line, to which a cutoff value of 0.5 is applied to obtain the resolution.

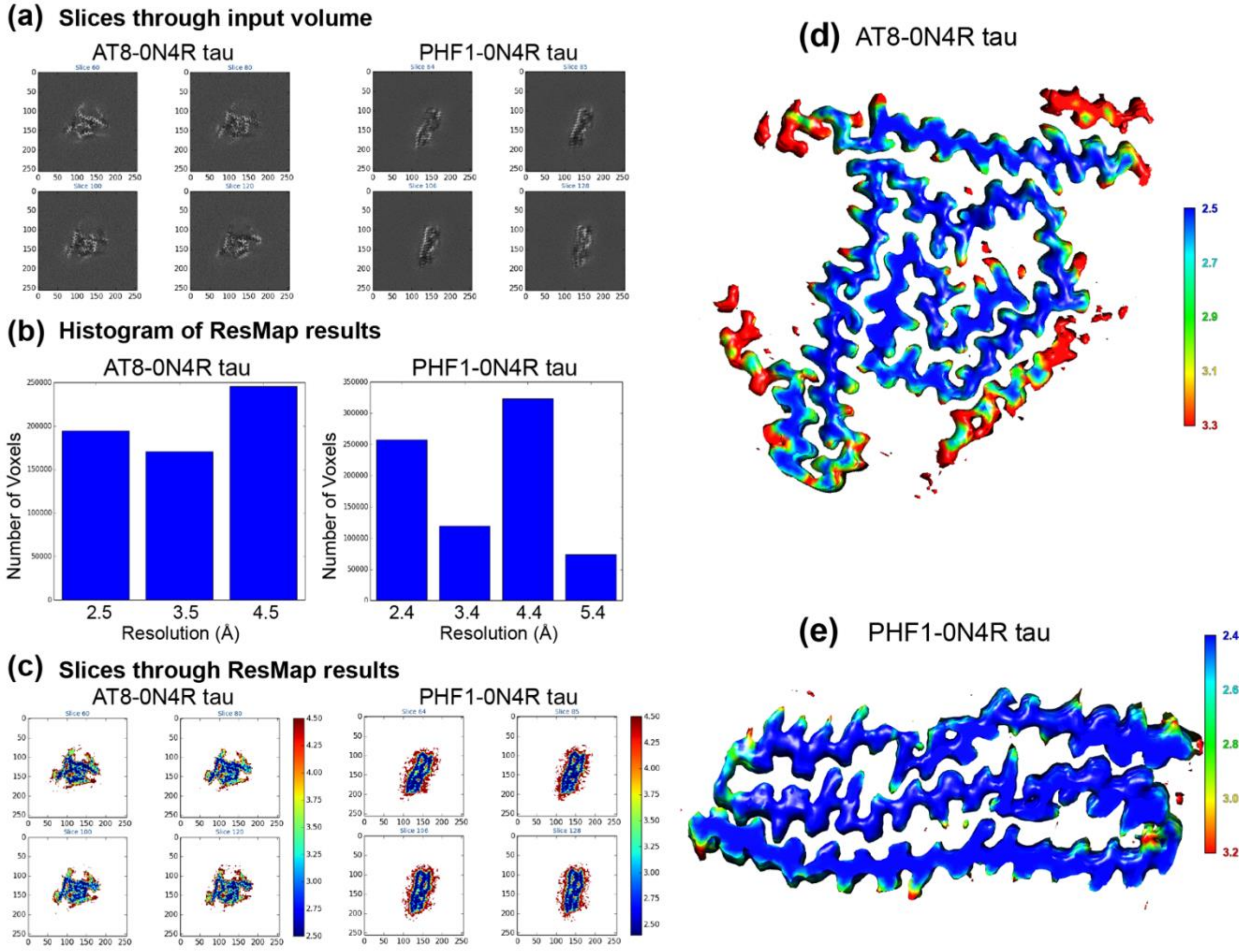


**Figure S11**. **Local resolution estimates of AT8- and PHF1-0N4R tau fibrils using the software ResMap**. (**a**) Density slices extracted by ResMap for the two fibrils. (**b**) Histograms of local resolution distribution. (**c**) Local resolution distribution of slices perpendicular to the fibril axis, highlight good local resolutions (blue, 2.5 Å to red, 4.5 Å) at the central region of fibrils. (**d**) 3D rendering of the local resolution analysis by ResMap of AT8-0N4R tau fibril. (**e**) 3D rendering of the local resolution of PHF1-0N4R tau fibrils.

**Table S1**. $^{13}C$ and $^{15}N$ chemical shifts of the AT8-0N4R tau fibrils. $^{13}C$ chemical shifts were referenced externally to the adamantane $CH_2$ chemical shift at 38.48 ppm on the tetramethylsilane scale. $^{15}N$ chemical shifts were referenced to the $^{15}N$ peak of N-acetylvaline at 122.00 ppm on the liquid ammonia scale.

| Resid | N | CO | Cα | Cβ | Cγ/γ1/γ# | Cγ2 | Cδ/δ1 | Cδ2/ε | Nγ2/δ2 |
|---|---|---|---|---|---|---|---|---|---|
| **V306** | | | 59.18 | | | | | | |
| **Q307** | 129.3 | 172.1 | 52.3 | 30.63 | 32.54 | | 177.3 | | 112.4 |
| **I308** | 126.8 | 171.9 | 57.7 | 38.49 | 25.24 | 15.06 | 11.64 | | |
| **V309** | 126.6 | 172 | 58.58 | 32.38 | 20.16 | 18.78 | | | |
| **Y310** | 128.6 | 171.9 | 57.86 | 38.7 | | | | 117.2 | |
| **K311** | 122.8 | 169.4 | 51.87 | 32.58 | | | | | |
| **P312** | 136.8 | 173.3 | 61.13 | 31.4 | 24.99 | | 48.77 | | |
| **V313** | 122.7 | 171.6 | 59.57 | 33.33 | 20.04 | 18.99 | | | |
| **D314** | 131.4 | 171.8 | 51.13 | 41.07 | 178 | | | | |
| **L315** | 125.1 | 173.6 | 51.37 | 42.93 | 25.51 | | | | |
| **S316** | 121.1 | | 54.34 | 63.32 | | | | | |
| **K317** | | 173 | 52.66 | | | | | | |
| **V318** | 127.4 | 172.8 | 58.95 | 33.52 | 19.01 | | | | |
| **T319** | 127.4 | 171.5 | 59.87 | 68.03 | 21.7 | | | | |
| **S320** | 119 | 173.8 | 52.82 | 62.67 | | | | | |
| **K321** | 123.3 | 173.8 | 56.62 | 27.62 | 25.44 | | 28.5 | 40.57 | |
| **C322** | 110.5 | 170.2 | 56.42 | 24.47 | | | | | |
| **G323** | 105.2 | 169.4 | 41.88 | | | | | | |
| **S324** | 118.2 | 171.9 | 53.86 | 61.15 | | | | | |
| **L325** | 129.9 | 174.3 | 52.52 | 41.83 | 27.49 | | 25.97 | 22.24 | |
| **G326** | 110.7 | 171.7 | 45.56 | | | | | | |
| **N327** | 118.3 | 172.7 | 51.19 | 38.94 | 173.5 | | | | 113.8 |
| **I328** | 120.9 | | 58.3 | 38.23 | 25.22 | 15.04 | 11.28 | | |
| **I354** | 119.1 | 176.2 | 55.23 | | 24.66 | 14.7 | | | |
| **G355** | 114.2 | 169.8 | 44.28 | | | | | | |
| **S356** | 110.1 | 171.5 | 54.74 | 63.89 | | | | | |
| **L357** | 120.3 | 175.3 | 52.71 | 43.41 | | | | | |
| **D358** | 120.4 | 172.4 | 51.39 | 45.51 | 180.1 | | | | |
| **N359** | 123 | 170.8 | 52.89 | 38.49 | 172.9 | | | | 108.2 |
| **I360** | 119.5 | 172.1 | 56.24 | 40.6 | 24.77 | 14.19 | 12.26 | | |
| **T361** | 123.4 | 170.5 | 59.51 | 65.9 | 21.76 | | | | |
| **H362** | 129.9 | 172 | 51.86 | 31.56 | | | | | |
| **V363** | 124.9 | 170.3 | 57.79 | 29.07 | 19.57 | | | | |
| **P364** | 135.6 | 175 | 60.4 | 29.97 | 25.6 | | 46.79 | | |
| **G365** | 105.3 | 169.2 | 42.5 | | | | | | |
| **G366** | 105.1 | 173.7 | 42.07 | | | | | | |
| **G367** | 106.2 | 172.5 | 45.1 | | | | | | |
| **N368** | 118.3 | | 51.19 | 38.97 | | | | | |
| **K369** | | 173 | | | | | | | |
| **K370** | 125.6 | 170.1 | 55.64 | 29.17 | 24.5 | | 25.38 | 40.51 | |
| **I371** | 125.7 | 174 | 55.57 | 34.99 | 24.34 | 16.92 | 10.91 | | |
| **E372** | 120.8 | 173.1 | 51.44 | 34.91 | 33.62 | | 180.2 | | |
| **T373** | 117.7 | 171.5 | 59.01 | 67.39 | 19.72 | | | | |
| **H374** | 125.4 | | 52.26 | 32.54 | 132.5 | | | 114.1 | |
| **K375** | | 172.2 | | | | | | | |
| **L376** | 128.2 | 173 | 53.42 | 43.1 | 28.86 | | 26.63 | 23.14 | |
| **T377** | 110.6 | 171.3 | 55.52 | 68.61 | 18.38 | | | | |
| **F378** | 120.7 | 172.9 | 54.08 | 37.92 | | | | | |
| **R379** | 131.4 | 172.1 | 51.86 | 31.82 | 25.69 | | 42.23 | | |
| **E380** | 120.7 | 172.1 | 54.32 | 22.85 | 31.89 | | 181.4 | | |

| | | | | | | | | | |
|---|---|---|---|---|---|---|---|---|---|
| **N381** | 111.7 | 173.4 | 49.31 | 41.56 | 175.3 | | | | 118.3 |
| **A382** | 121.3 | 173.4 | 49.83 | 12.63 | | | | | |
| **K383** | 119.4 | 172.2 | 52.02 | 34.58 | 22.36 | | 28.47 | 39.87 | |
| **A384** | 123.7 | 173.9 | 48.04 | 19.33 | | | | | |
| **K385** | 127.8 | 172.6 | 52.83 | 32.56 | 23.67 | | 27.54 | 39.79 | |
| **T386** | 129 | 170.8 | 59.77 | 66.4 | 20.04 | | | | |
| **D387** | 127.7 | 172.8 | 49.17 | 43.24 | 177.5 | | | | |
| **H388** | 116.9 | 174.6 | 55.74 | 25.77 | 130.6 | | 117.3 | 135 | |
| **G389** | 103.2 | 170.2 | 44.59 | | | | | | |
| **A390** | 122.7 | 172.4 | 50.13 | 19.17 | | | | | |
| **E391** | 118.5 | 172.5 | 51.93 | 31.35 | | | 179.8 | | |
| **I392** | 124.7 | 172.1 | 57.44 | 38.8 | 25.5 | 14.45 | 11.61 | | |
| **V393** | 128.8 | 171.7 | 59.52 | 30.89 | 18.73 | | | | |
| **Y394** | 129.5 | 173.2 | 54.92 | 39.41 | | | | | |
| **K395** | 129.1 | 171.1 | 55.04 | 29.82 | 25.96 | | 29.28 | 41.53 | |
| **S396** | 109.3 | 167.6 | 51.94 | 61.55 | | | | | |
| **P397** | 129.7 | 171.5 | 59.64 | 32.08 | 24.17 | | 48.31 | | |
| **V398** | 117.7 | 173 | 58.74 | 31.11 | 18.75 | | | | |
| **V399** | 126.5 | 173.1 | 58.84 | 33 | 18.75 | | | | |
| **S400** | 122.6 | 175 | 54.39 | 62.67 | | | | | |
| **G401** | 111.8 | 167.8 | 42.6 | | | | | | |
| **D402** | 117.5 | 174.9 | 51.05 | 38.25 | 178.3 | | | | |
| **T403** | 124.2 | 171.7 | 59.32 | 66.2 | 18.68 | | | | |
| **S404** | 120.8 | 168.6 | 51.94 | 60.56 | | | | | |
| **P405** | 137.1 | | 58.23 | 29.59 | 25.63 | | 45.44 | | |
| **S409** | | 169.8 | 54.17 | | | | | | |
| **N410** | 127 | 172.5 | 51.01 | 40.51 | 172 | | | | 115.3 |
| **V411** | 125.5 | 172.7 | 58.47 | 35.63 | 19.87 | 18.67 | | | |
| **S412** | 120.6 | 171.2 | 54.43 | 62.54 | | | | | |
| **S413** | 124.5 | 172.1 | 53.19 | 62.38 | | | | | |
| **T414** | 127.2 | 172.4 | 58.51 | 68.79 | 21.43 | | | | |
| **G415** | 117.7 | 171.4 | 47.19 | | | | | | |
| **S416** | 113.8 | 170.4 | 54.26 | 64.12 | | | | | |
| **I417** | 122 | 172.1 | 58.85 | 39.61 | 26.84 | 15.21 | 11.64 | | |
| ***D418*** | 130.7 | 171.7 | 50.8 | 40.41 | 177.1 | | | | |
| **M419** | 124.9 | 172 | 52.63 | 34.51 | | | | | |
| **V420** | 124.9 | 173.9 | 58.33 | 32.2 | 18.78 | | | | |
| **D421** | 128.1 | 170.8 | 51.36 | 34.11 | 178.3 | | | | |
| **S422** | 110.7 | 166.4 | 52.78 | 63 | | | | | |
| **P423** | 127.3 | 171 | 59.15 | 32.16 | 26.36 | | 47.8 | | |
| **Q424** | 116.1 | 171.9 | 51.94 | 30.79 | 30.72 | | 175.8 | | 108.2 |
| **L425** | 125 | 171.6 | 52.67 | 41.78 | 27.67 | | 26.7 | 22.26 | |
| **A426** | 125.1 | 173.4 | 47.6 | 19.97 | | | | | |
| **T427** | 119.9 | 171.6 | 57.31 | 69.85 | 20.94 | | | | |
| **L428** | 132.6 | 172.6 | 52.86 | 37.82 | 27.17 | | 23.02 | | |
| **A429** | 115.2 | 174.3 | 47.86 | 22.51 | | | | | |
| **E431** | | 170.8 | | | | | | | |
| **V432** | 124.1 | 172.8 | 58.73 | 31.35 | 18.4 | | | | |
| **S433** | 118.8 | 170.7 | 53.44 | 64.19 | | | | | |
| **A434** | 120 | 176.5 | 51.67 | 18.13 | | | | | |
| **S435** | 110.9 | 171.9 | 54.94 | 62.18 | | | | | |
| **L436** | 135 | 172.1 | 52.72 | 42.99 | 26.87 | | 23.36 | | |
| **A437** | 128.9 | 173.3 | 48.01 | 18.64 | | | | | |
| **K438** | 129.1 | 171.9 | 50.86 | 34.69 | 23.72 | | 26.12 | | |
| **Q439** | 123.9 | 173.2 | 51.63 | 31.03 | 31.29 | | 175.7 | | 111.5 |
| **G440** | 109.6 | | 43.05 | | | | | | |

**Table S2**. $^{13}$C and $^{15}$N chemical shifts of the PHF1-0N4R tau fibrils. $^{13}$C chemical shifts were referenced externally to the adamantane $CH_2$ chemical shift at 38.48 ppm on the tetramethylsilane scale. $^{15}$N chemical shifts were referenced to the $^{15}$N peak of N-acetylvaline at 122.00 ppm on the liquid ammonia scale.

| Resid | N | C | Ca | Cb | Cg/ Cg1 | Cg2 | Cd/Cd1 | Cd2 | Ce | Cz | Nd | Ne | Nz |
|---|---|---|---|---|---|---|---|---|---|---|---|---|---|
| **V275** | 126 | 172.5 | 59.42 | 33.43 | 19.95 | 18.01 | | | | | | | |
| **Q276** | 129.4 | 171.8 | 52.48 | 31.65 | 33.04 | | 177.8 | | | | | | |
| **I277** | 127.3 | 171.7 | 58.51 | 39.03 | 25.57 | 15.63 | 11.3 | | | | | | |
| **I278** | 127.8 | 171.5 | 57.91 | 38.51 | 25.64 | 15.61 | 11.31 | | | | | | |
| **N279** | 128.1 | 171.6 | 50.08 | 40.55 | 173.7 | | | | | | 113.2 | | |
| **K280** | 125.9 | | 52.81 | 33.1 | | | | | | | | | |
| **V287** | 128.4 | 173.8 | 59.34 | 31.98 | 20.21 | 17.63 | | | | | | | |
| **Q288** | 119.8 | 173.7 | 51.55 | 36.73 | 33.58 | | 178.7 | | | | | 116.3 | |
| **S289** | 112.3 | 172.7 | 52.34 | 63.28 | | | | | | | | | |
| **K290** | 127.5 | 173.3 | 56.39 | 29.86 | 27.15 | | 29.85 | | 40.62 | | | | |
| **C291** | 115.7 | 172.8 | 56.29 | 28.12 | | | | | | | | | |
| **G292** | 115.8 | 170.9 | 46.62 | | | | | | | | | | |
| **S293** | 115.3 | 172.6 | 54.11 | 64.7 | | | | | | | | | |
| **K294** | 124.7 | 172.4 | 54.24 | 34.43 | | | | | | | | | |
| **D295** | 128.3 | 172.5 | 51.66 | 43.22 | 181 | | | | | | | | |
| **N296** | 118.7 | 171.9 | 51 | 43.11 | 175.4 | | | | | | 117.4 | | |
| **I297** | 122.3 | 173.5 | 59.16 | 38.3 | 25.36 | 15.51 | 13.96 | | | | | | |
| **K298** | 119.7 | 171.6 | 54.92 | | | | | | | | | | |
| **V306** | 128.9 | 170.8 | 59.29 | 32.53 | 18.45 | | | | | | | | |
| **Q307** | 131.2 | 172.3 | 52.02 | 32.5 | 32.21 | | 175.9 | | | | | 111.3 | |
| **I308** | 125 | 172.9 | 57.32 | 40.92 | 25.48 | 16.01 | 12.08 | | | | | | |
| **V309** | 128.5 | 171.7 | 59.61 | 31.14 | 20.15 | 19.11 | | | | | | | |
| **Y310** | 132 | 169.3 | 56.2 | 40.74 | 123.7 | | 129.9 | | 115.4 | 157.9 | | | |
| **K311** | 130.3 | 169 | 55.23 | 32.17 | 25.45 | | 29.92 | | 41.33 | | | | |
| **P312** | 125.7 | 174.7 | 60.58 | 30.25 | 26.89 | | 46.88 | | | | | | |
| **V313** | 119.7 | 169.1 | 61.71 | 25.88 | 21.94 | 20.31 | | | | | | | |
| **D314** | 124.6 | 174.8 | 51.09 | 39.04 | 180 | | | | | | | | |
| **L315** | 119 | 175.8 | 51.13 | 43.32 | 25.43 | | | | | | | | |
| **S316** | 115.5 | 170.8 | 53.92 | 63.42 | | | | | | | | | |
| **K317** | 130.2 | 172.1 | 52.87 | 35.55 | 23.79 | | 28.76 | | 39.94 | | | | |
| **V318** | 126 | 173.5 | 58.58 | 33.91 | 19.33 | | | | | | | | |
| **T319** | 128.4 | 171.1 | 60.12 | 67.52 | 22.03 | | | | | | | | |
| **S320** | 121.3 | 172.7 | 53.14 | 63.02 | | | | | | | | | |
| **K321** | 127.5 | 173.2 | 56.04 | 28.19 | 25.69 | | 28.99 | | 41.57 | | | | |
| **C322** | 110.7 | 171.8 | 55.63 | 28.92 | | | | | | | | | |
| **G323** | 111.1 | 169.7 | 42.99 | | | | | | | | | | |
| **S324** | 121.9 | 173.1 | 53.21 | 60.24 | | | | | | | | | |
| **L325** | 128.3 | 176.5 | 53.89 | 43.19 | 28.64 | | 25.91 | 22.03 | | | | | |
| **G326** | 109.2 | 172.1 | 45.72 | | | | | | | | | | |
| **N327** | 119.7 | 172.3 | 51.96 | 39.45 | 173.9 | | | | | | 119.9 | | |
| **I328** | 120.2 | 172 | 58.96 | 39.93 | 25.36 | 15.58 | 12 | | | | | | |
| **H329** | 126.6 | | 52.79 | | | | | | 136.5 | | | | |
| **H330** | | 170.8 | | | | | 116.2 | | 136.4 | | | | |
| **K331** | 126.4 | 168.7 | 51.6 | 31.32 | 22.73 | | | | | | | | |
| **P332** | 134.8 | 174.9 | 60.88 | 30.74 | 26.46 | | 46.84 | | | | | | |
| **G333** | 110.5 | 171.6 | 44.15 | | | | | | | | | | |
| **G334** | 116.6 | 174.5 | 43.55 | | | | | | | | | | |
| **G335** | 107.1 | 170.7 | 45.42 | | | | | | | | | | |
| **Q336** | 113.4 | 171 | 52.34 | 31.71 | | | 176.1 | | | | | 111.4 | |
| **V337** | 119 | 173.6 | 59.38 | 33.91 | 19.52 | | | | | | | | |
| **E338** | 132.9 | 171.4 | 53.1 | 33.41 | 36.13 | | 181.4 | | | | | | |

| | | | | | | | | | | |
|---|---|---|---|---|---|---|---|---|---|---|
| **V339** | 128.7 | - | 58.78 | 33.14 | 19.62 | | | | | |
| **E342** | | 174.2 | | | | | | | | |
| **K343** | 125.9 | 172.1 | 53.38 | 34.57 | 23.95 | | 28.15 | | 40.22 | 33.94 |
| **L344** | 129.7 | 171.7 | 53.45 | 41.16 | 25.82 | | 28.25 | 22.01 | | |
| **D345** | 125.9 | 172.4 | 50.35 | 40.3 | 178.6 | | | | | |
| **F346** | 123.6 | 175 | 49.76 | 36.86 | 137.2 | | | | | |
| **K347** | 125.2 | 172.6 | 54.28 | 35.69 | 24.42 | | 28.68 | | 41.29 | 34.34 |
| **D348** | 124.8 | 174.1 | 50.75 | 42.35 | 179.1 | | | | | |
| **R349** | 121.8 | 172.7 | 52.97 | 32.69 | 25.7 | | 40.22 | | | |
| **V350** | 125.9 | 172.4 | 59.14 | 33.63 | 19.52 | | | | | |
| **Q351** | 127.9 | 172.3 | 52.67 | 31.54 | 33.35 | | 178.3 | | | |
| **S352** | 116.9 | 171.9 | 54.07 | 63.54 | | | | | | |
| **K353** | 131.8 | 172.8 | 54.05 | 34.93 | 23.96 | | 28.18 | | 40.31 | |
| **I354** | 129.5 | 173.7 | 59.32 | 40.3 | 27.27 | 14.55 | 11.37 | | | |
| **G355** | 113.9 | 169.8 | 43.57 | | | | | | | |
| **S356** | 113 | 170.8 | 56.97 | 62.65 | | | | | | |
| **L357** | 123.4 | | 53.94 | | | | | | | |

**Table S3**. 2D $^{13}C$-$^{13}C$ TOCSY intensity ratios of several well-resolved residue types in AT8, PHF1 and AT8/PHF1 0N4R tau fibrils. Observed intensity ratios are compared with the predicted intensity ratios estimated based on the distribution of these amino acid types in the protein sequence and the proposed fuzzy coat dynamics shown in Figure 5d.

| Intensity ratio | AT8-0N4R tau | | PHF1-0N4R tau | | AT8/PHF1 0N4R tau | |
|---|---|---|---|---|---|---|
| | Observed | Predicted | Observed | Predicted | Observed | Predicted |
| Glu-Pro / Glu | 1.14 | 1.00 | 0.53 | 0.40 | 0.53 | 0.50 |
| Ser-Pro / Ser | 0.12 | 0.10 | 0.19 | 0.18 | 0.09 | 0.13 |
| Thr-Pro /Thr | 0.74 | 1.00 | 0.79 | 1.10 | 0.64 | 0.70 |
| Ala-Pro /Ala | 0.17 | 0.21 | 0.10 | 0.21 | 0.078 | 0.18 |

**Table S4**. Solid-state NMR experimental parameters for three phospho-mimetic tau fibrils. All experiments were recorded on an 18.8 T spectrometer (800 MHz $^1$H frequency) using a BlackFox 3.2 mm E-Free HCN probe. Reported temperatures are estimated sample temperatures based on external calibration of probe thermocouple and the water $^1$H chemical shift.

| Experiment | NMR Parameters | Expt. Time |
|---|---|---|
| ***AT8-0N4R tau*** | | |
| 2D CC CORD | $T_{sample}$ = 273 K; $\nu_{MAS}$ = 10.5 kHz, ns = 48, $\tau_{rd}$ = 1.6 s, $t_{1,max}$ = 9.9 ms; $t_{1,inc}$ = 24.8 μs; $\tau_{dwell}$ = 6 μs; $\tau_{acq}$ = 15.4 ms; $\tau_{HC}$ = 70 μs; $\tau_{CORD}$ = 23 ms; $\nu_{1Hacq}$ = 83 kHz TPPM | 17.5 hr |
| 2D NCA SPEC-CP | $T_{sample}$ = 273 K; $\nu_{MAS}$ = 10.5 kHz, ns = 128, $\tau_{rd}$ = 1.7 s, $t_{1,max}$ = 9.5 ms; $t_{1,inc}$ = 95.2 μs; $\tau_{dwell}$ = 6 μs; $\tau_{acq}$ = 15.4 ms; $\tau_{HN}$ = 0.8 ms; $\tau_{NC}$ = 4.5 ms; $\nu_{1HspecificCP}$ = 83 kHz CW; $\nu_{1Hacq}$ = 83 kHz TPPM | 12.5 hr |
| 2D NCACB SPECCP-DREAM | $T_{sample}$ = 278 K; $\nu_{MAS}$ = 14 kHz, ns = 88, $\tau_{rd}$ = 1.7 s, $t_{1,max}$ = 14.3 ms; $t_{1,inc}$ = 71.4 μs; $\tau_{dwell}$ = 6 μs; $\tau_{acq}$ = 15.4 ms; $\tau_{HN}$ = 1 ms; $\tau_{NC}$ = 4.5 ms; $\tau_{CC}$ = 2.5 ms; $fq_{13Cdream}$ = 55 ppm; $\nu_{1HspecificCP}$ = 83 kHz CW; $\nu_{1Hdream}$ = 83 kHz CW; $\nu_{1Hacq}$ = 83 kHz TPPM | 16 hr |
| 2D $H^N$ rINEPT | $T_{sample}$ = 273 K; $\nu_{MAS}$ = 10.5 kHz, ns = 16, $\tau_{rd}$ = 1.5 s, $t_{1,max}$ = 21.9 ms; $t_{1,inc}$ = 124.9 μs; $\tau_{dwell}$ = 20 μs; $\tau_{acq}$ = 81.9 ms; $\nu_{1Hacq}$ = 18 kHz TPPM, $\nu_{13Cacq}$ = 10 kHz GARP | 2.5 hr |
| 2D CC TOCSY<br>High temperature | $T_{sample}$ = 298 K; $\nu_{MAS}$ = 14 kHz, ns = 32, $\tau_{rd}$ = 1.8 s, $t_{1,max}$ = 7.5 ms; $t_{1,inc}$ = 24.9 μs; $\tau_{dwell}$ = 6 μs; $\tau_{acq}$ = 24.6 ms; $\tau_{TOCSY}$ = 12.2 ms; $\nu_{13C\ TOCSY}$ = 36 kHz DIPSI-3 at 50 ppm; $\nu_{1H\ TOCSY}$ = 0 kHz; $\nu_{1Hacq}$ = 50 kHz TPPM | 10 hr |
| 3D NCACX SPECCP-CORD | $T_{sample}$ = 278 K; $\nu_{MAS}$ = 14 kHz, ns = 48, $\tau_{rd}$ = 1.7 s, $t_{1,max}$ = 6.9 ms; $t_{1,inc}$ = 142.9 μs; $t_{2,max}$ = 4.3 ms; $t_{2,inc}$ = 142.9 μs; $\tau_{dwell}$ = 6 μs; $\tau_{acq}$ = 15.4 ms; $\tau_{HN}$ = 1 ms; $\tau_{NC}$ = 4.5 ms; $\nu_{1HspecificCP}$ = 83 kHz CW; $\tau_{CORD}$ = 82 ms; $\nu_{1Hacq}$ = 83 kHz TPPM | 140 hr |
| 3D NCOCX SPECCP-CORD | $T_{sample}$ = 278 K; $\nu_{MAS}$ = 14 kHz, ns = 48, $\tau_{rd}$ = 1.65 s, $t_{1,max}$ = 6.9 ms; $t_{1,inc}$ = 142.9 μs; $t_{2,max}$ = 4.3 ms; $t_{2,inc}$ = 142.9 μs; $\tau_{dwell}$ = 6 μs; $\tau_{acq}$ = 15.4 ms; $\tau_{HN}$ = 1 ms; $\tau_{NC}$ = 4.5 ms; $\nu_{1HspecificCP}$ = 83 kHz CW; $\tau_{CORD}$ = 82 ms; $\nu_{1Hacq}$ = 83 kHz TPPM | 136 hr |
| 3D CONCA SPECCP-SPECCP | $T_{sample}$ = 278 K; $\nu_{MAS}$ = 14 kHz, ns = 32, $\tau_{rd}$ = 1.6 s; $t_{1,max}$ = 4.9 ms; $t_{1,inc}$ = 214.3 μs; $t_{2,max}$ = 6.9 ms; $t_{2,inc}$ = 142.9 μs; $\tau_{dwell}$ = 6 μs; $\tau_{acq}$ = 15.4 ms; $\tau_{HC}$ = 1 ms; $\tau_{CN}$ = 4.5 ms; $\tau_{NC}$ = 4.5 ms; $\nu_{1HspecificCP}$ = 83 kHz CW; $\nu_{1Hacq}$ = 83 kHz TPPM | 64 hr |
| 3D CONCACB SPECCP-SPECCP-DREAM | $T_{sample}$ = 278 K; $\nu_{MAS}$ = 14 kHz, ns = 64, $\tau_{rd}$ = 1.6 s; $t_{1,max}$ = 4.9 ms; $t_{1,inc}$ = 214.3 μs; $t_{2,max}$ = 6.9 ms; $t_{2,inc}$ = 142.9 μs; $\tau_{dwell}$ = 6 μs; $\tau_{acq}$ = 15.4 ms; $\tau_{HC}$ = 1 ms; $\tau_{CN}$ = 4.5 ms; $\tau_{NC}$ = 4.5 ms; $\tau_{CC}$ = 2.5 ms at 55 ppm; $\nu_{1HspecificCP}$ = 83 kHz CW; $\nu_{1Hdream}$ = 83 kHz CW; $\nu_{1Hacq}$ = 83 kHz TPPM | 128 hr |
| 3D CAN(CO)CA SPECCP-SPECCP-BSHCP | $T_{sample}$ = 278 K; $\nu_{MAS}$ = 14 kHz, ns = 64, $\tau_{rd}$ = 1.6 s; $t_{1,max}$ = 5 ms; $t_{1,inc}$ = 142.9 μs; $t_{2,max}$ = 7.5 ms; $t_{2,inc}$ = 214.3 μs; $\tau_{dwell}$ = 6 μs; $\tau_{acq}$ = 13.5 ms; $\tau_{HC}$ = 170 μs; $\tau_{CN}$ = 4.5 ms; $\tau_{NC}$ = 4.5 ms; $\tau_{CC}$ = 3.5 ms; $fq_{13CbshCP}$ = 90 ppm; $\nu_{1HspecificCP}$ = 83 kHz CW; $\nu_{1HbshCP}$ = 83 kHz CW; $\nu_{1Hacq}$ = 83 kHz TPPM | 144 hr |
| 2D Water-edited NCACB SPECCP-DREAM | $T_{sample}$ = 278 K; $\nu_{MAS}$ = 14 kHz, ns = 896, $\tau_{rd}$ = 1.7 s, $t_{1,max}$ = 7.1 ms; $t_{1,inc}$ = 71.4 μs; $\tau_{dwell}$ = 6 μs; $\tau_{acq}$ = 15.4 ms; $\tau_{1Hsel-echo-pulse}$ = 0.95 ms; $\tau_{1Hsel-echo}$ = 2x0.095 ms; $\tau_{1HSD}$ = 4 ms; $\tau_{HN}$ = 1 ms; $\tau_{NC}$ = 4.5 ms; $\tau_{CC}$ = 2.5 ms; $fq_{13Cdream}$ = 55 ppm; $\nu_{1HspecificCP}$ = 83 kHz CW; $\nu_{1Hdream}$ = 83 kHz CW; $\nu_{1Hacq}$ = 83 kHz TPPM | 87 hr |
| ***PHF1-0N4R tau*** | | |

| | | |
|---|---|---|
| 2D CC CORD | $T_{sample}$ = 270 K; $\nu_{MAS}$ = 10.5 kHz, ns = 48, $\tau_{rd}$ = 1.7 s, $t_{1,max}$ = 9.9 ms; $t_{1,inc}$ = 24.8 μs; $\tau_{dwell}$ = 6 μs; $\tau_{acq}$ = 15.4 ms; $\tau_{HC}$ = 70 μs; $\tau_{CORD}$ = 23 ms; $\nu_{1Hacq}$ = 83 kHz TPPM | 18.5 hr |
| 2D CC DREAM | $T_{sample}$ = 273 K; $\nu_{MAS}$ = 14 kHz, ns = 64, $\tau_{rd}$ = 1.7 s, $t_{1,max}$ = 6.2 ms; $t_{1,inc}$ = 24.8 μs; $\tau_{dwell}$ = 6 μs; $\tau_{acq}$ = 15.4 ms; $\tau_{HC}$ = 70 μs; $\tau_{CC}$ = 2.5 ms; $fq_{13Cdream}$ = 40 ppm; $\nu_{1Hdream}$ = 83 kHz TPPM; $\nu_{1Hacq}$ = 83 kHz TPPM | 15 hr |
| 2D NCA SPEC-CP | $T_{sample}$ = 270 K; $\nu_{MAS}$ = 10.5 kHz, ns = 144, $\tau_{rd}$ = 1.7 s, $t_{1,max}$ = 11.9 ms; $t_{1,inc}$ = 95.2 μs; $\tau_{dwell}$ = 6 μs; $\tau_{acq}$ = 15.4 ms; $\tau_{HN}$ = 1 ms; $\tau_{NC}$ = 5 ms; $\nu_{1HspecificCP}$ = 83 kHz CW; $\nu_{1Hacq}$ = 83 kHz TPPM | 17.5 hr |
| 2D NCACB SPECCP-DREAM | $T_{sample}$ = 273 K; $\nu_{MAS}$ = 14 kHz, ns = 128, $\tau_{rd}$ = 1.7 s, $t_{1,max}$ = 14.3 ms; $t_{1,inc}$ = 71.4 μs; $\tau_{dwell}$ = 6 μs; $\tau_{acq}$ = 15.4 ms; $\tau_{HN}$ = 1 ms; $\tau_{NC}$ = 3.5 ms; $\tau_{CC}$ = 2.5 ms; $fq_{13Cdream}$ = 55 ppm; $\nu_{1HspecificCP}$ = 83 kHz CW; $\nu_{1Hdream}$ = 83 kHz CW; $\nu_{1Hacq}$ = 83 kHz TPPM | 25 hr |
| 2D HN rINEPT | $T_{sample}$ = 273 K; $\nu_{MAS}$ = 14 kHz, ns = 16, $\tau_{rd}$ = 2 s, $t_{1,max}$ = 31.2 ms; $t_{1,inc}$ = 124.9 μs; $\tau_{dwell}$ = 20 μs; $\tau_{acq}$ = 81.9 ms; $\nu_{1Hacq}$ = 18 kHz TPPM, $\nu_{13Cacq}$ = 10 kHz GARP | 5 hr |
| 2D CC TOCSY<br>High temperature | $T_{sample}$ = 298 K; $\nu_{MAS}$ = 14 kHz, ns = 32, $\tau_{rd}$ = 2 s, $t_{1,max}$ = 9.1 ms; $t_{1,inc}$ = 24.9 μs; $\tau_{dwell}$ = 6 μs; $\tau_{acq}$ = 24.6 ms; $\tau_{TOCSY}$ = 12.2 ms; $\nu_{13C\ TOCSY}$ = 36 kHz DIPSI-3 at 50 ppm; $\nu_{1H\ TOCSY}$ = 0 kHz; $\nu_{1Hacq}$ = 50 kHz TPPM | 13.5 hr |
| 3D NCACX SPECCP-CORD | $T_{sample}$ = 273 K; $\nu_{MAS}$ = 14 kHz, ns = 48, $\tau_{rd}$ = 1.7 s, $t_{1,max}$ = 6.4 ms; $t_{1,inc}$ = 142.9 μs; $t_{2,max}$ = 4.8 ms; $t_{2,inc}$ = 142.9 μs; $\tau_{dwell}$ = 6 μs; $\tau_{acq}$ = 15.4 ms; $\tau_{HN}$ = 1 ms; $\tau_{NC}$ = 4.5 ms; $\nu_{1HspecificCP}$ = 83 kHz CW; $\tau_{CORD}$ = 82 ms; $\nu_{1Hacq}$ = 83 kHz TPPM | 149 hr |
| 3D NCOCX SPECCP-CORD | $T_{sample}$ = 273 K; $\nu_{MAS}$ = 14 kHz, ns = 48, $\tau_{rd}$ = 1.7 s, $t_{1,max}$ = 6.4 ms; $t_{1,inc}$ = 142.9 μs; $t_{2,max}$ = 4.6 ms; $t_{2,inc}$ = 142.9 μs; $\tau_{dwell}$ = 6 μs; $\tau_{acq}$ = 15.4 ms; $\tau_{HN}$ = 1 ms; $\tau_{NC}$ = 4.5 ms; $\nu_{1HspecificCP}$ = 83 kHz CW; $\tau_{CORD}$ = 82 ms; $\nu_{1Hacq}$ = 83 kHz TPPM | 140 hr |
| 3D CONCA SPECCP-SPECCP | $T_{sample}$ = 273 K; $\nu_{MAS}$ = 14 kHz, ns = 32, $\tau_{rd}$ = 1.7 s; $t_{1,max}$ = 4.9 ms; $t_{1,inc}$ = 214.3 μs; $t_{2,max}$ = 6.9 ms; $t_{2,inc}$ = 142.9 μs; $\tau_{dwell}$ = 6 μs; $\tau_{acq}$ = 15.4 ms; $\tau_{HC}$ = 1 ms; $\tau_{CN}$ = 4.5 ms; $\tau_{NC}$ = 5 ms; $\nu_{1HspecificCP}$ = 83 kHz CW; $\nu_{1Hacq}$ = 83 kHz TPPM | 68 hr |
| 2D Water-edited NCACB SPECCP-DREAM | $T_{sample}$ = 273 K; $\nu_{MAS}$ = 14 kHz, ns = 864, $\tau_{rd}$ = 1.7 s, $t_{1,max}$ = 7.1 ms; $t_{1,inc}$ = 71.4 μs; $\tau_{dwell}$ = 6 μs; $\tau_{acq}$ = 15.4 ms; $\tau_{1Hsel\text{-}echo\text{-}pulse}$ = 0.95 ms; $\tau_{1Hsel\text{-}echo}$ = 2x0.095 ms; $\tau_{1HSD}$ = 4 ms; $\tau_{HN}$ = 1 ms; $\tau_{NC}$ = 3.5 ms; $\tau_{CC}$ = 2.5 ms; $fq_{13Cdream}$ = 55 ppm; $\nu_{1HspecificCP}$ = 83 kHz CW; $\nu_{1Hdream}$ = 83 kHz CW; $\nu_{1Hacq}$ = 83 kHz TPPM | 84 hr |
| 2D Water-edited CC DREAM | $T_{sample}$ = 273 K; $\nu_{MAS}$ = 14 kHz, ns = 224, $\tau_{rd}$ = 1.7 s, $t_{1,max}$ = 6.2 ms; $t_{1,inc}$ = 24.8 μs; $\tau_{dwell}$ = 6 μs; $\tau_{acq}$ = 15.4 ms; $\tau_{1Hsel\text{-}echo\text{-}pulse}$ = 0.95 ms; $\tau_{1Hsel\text{-}echo}$ = 2x0.095 ms; $\tau_{1HSD}$ = 4 ms; $\tau_{HC}$ = 70 μs; $\tau_{CC}$ = 2.5 ms; $fq_{13Cdream}$ = 40 ppm; $\nu_{1Hdream}$ = 83 kHz TPPM; $\nu_{1Hacq}$ = 83 kHz TPPM | 54 hr |
| ***AT8/PHF1 0N4R tau*** | | |
| 2D CC CORD | $T_{sample}$ = 270 K; $\nu_{MAS}$ = 10.5 kHz, ns = 48, $\tau_{rd}$ = 1.7 s, $t_{1,max}$ = 9.9 ms; $t_{1,inc}$ = 24.8 μs; $\tau_{dwell}$ = 6 μs; $\tau_{acq}$ = 15.4 ms; $\tau_{HC}$ = 70 μs; $\tau_{CORD}$ = 23 ms; $\nu_{1Hacq}$ = 83 kHz TPPM | 18.5 hr |
| 2D CC DREAM | $T_{sample}$ = 273 K; $\nu_{MAS}$ = 14 kHz, ns = 64, $\tau_{rd}$ = 1.7 s, $t_{1,max}$ = 7.5 ms; $t_{1,inc}$ = 24.9 μs; $\tau_{dwell}$ = 6 μs; $\tau_{acq}$ = 15.4 ms; $\tau_{HC}$ = 70 μs; $\tau_{CC}$ = 2.5 ms; $fq_{13Cdream}$ = 40 ppm; $\nu_{1Hdream}$ = 83 kHz TPPM; $\nu_{1Hacq}$ = 71 kHz TPPM | 15 hr |

| | | |
|---|---|---|
| 2D NCA SPEC-CP | $T_{sample}$ = 270 K; $\nu_{MAS}$ = 10.5 kHz, ns = 144, $\tau_{rd}$ = 1.7 s, $t_{1,max}$ = 12.4 ms; $t_{1,inc}$ = 95.2 μs; $\tau_{dwell}$ = 6 μs; $\tau_{acq}$ = 15.4 ms; $\tau_{HN}$ = 1 ms; $\tau_{NC}$ = 5 ms; $\nu_{1HspecificCP}$ = 83 kHz CW; $\nu_{1Hacq}$ = 83 kHz TPPM | 18 hr |
| 2D HN rINEPT | $T_{sample}$ = 273 K; $\nu_{MAS}$ = 14 kHz, ns = 64, $\tau_{rd}$ = 2 s, $t_{1,max}$ = 31.2 ms; $t_{1,inc}$ = 125.0 μs; $\tau_{dwell}$ = 20 μs; $\tau_{acq}$ = 81.9 ms; $\nu_{1Hacq}$ = 18 kHz TPPM, $\nu_{13Cacq}$ = 10 kHz GARP | 19 hr |
| 2D CC TOCSY<br>High temperature | $T_{sample}$ = 298 K; $\nu_{MAS}$ = 14 kHz, ns = 64, $\tau_{rd}$ = 2 s, $t_{1,max}$ = 9.1 ms; $t_{1,inc}$ = 24.9 μs; $\tau_{dwell}$ = 6 μs; $\tau_{acq}$ = 24.6 ms; $\tau_{TOCSY}$ = 12.2 ms; $\nu_{13C\ TOCSY}$ = 36 kHz DIPSI-3 at 50 ppm; $\nu_{1H\ TOCSY}$ = 0 kHz; $\nu_{1Hacq}$ = 50 kHz TPPM | 27 hr |

$T_{sample}$ = estimated sample temperature based on water chemical shift; $\nu_{MAS}$ = magic angle spinning frequency; ns = number of scans per free induction decay; $\tau_{rd}$ = recycle delay; $t_{1,max}$ = maximum $t_1$ evolution time; $t_{1,inc}$ = $t_1$ increment; $t_{2,max}$ = maximum $t_2$ evolution time; $t_{2,inc}$ = $t_2$ increment; $\tau_{dwell}$ = dwell-time in the direct dimension; $\tau_{acq}$ = maximum acquisition time in the direct dimension; $\tau_{HC}$ = $^1$H-$^{13}$C cross polarization contact time; $\tau_{CORD}$ = $^{13}$C-$^{13}$C mixing time using CORD; $\tau_{HN}$ = $^1$H-$^{15}$N cross polarization contact time; $\tau_{NC}$ = $^{15}$N-$^{13}$C specific cross polarization contact time; $\tau_{CC}$ = 13C-13C DREAM or $^{BSH}$CP cross polarization time; $\nu_{1Hacq}$ = $^1$H rf field strength for decoupling during acquisition; $\nu_{13C\ TOCSY}$ = $^{13}$C rf nutation frequency and rf carrier frequency for TOCSY transfer; $fq_{13Cdream}$ = $^{13}$C rf carrier frequency during DREAM spin-lock; $\tau_{1Hsel\text{-}echo\text{-}pulse}$ = $^1$H 180° Gauss5% pulse length for selective echo; $\tau_{1Hsel\text{-}echo}$ = additional $T_2$ time outside pulse in selective echo; $\tau_{1HSD}$ = $^1$H-$^1$H spin diffusion mixing time.
